# EEG-Based Spectral Analysis Showing Brainwave Changes Related to Modulating Progressive Fatigue During a Prolonged Intermittent Motor Task

**DOI:** 10.1101/2021.09.08.458591

**Authors:** Easter S. Suviseshamuthu, Vikram Shenoy Handiru, Didier Allexandre, Armand Hoxha, Soha Saleh, Guang H. Yue

## Abstract

Repeatedly performing a submaximal motor task for a prolonged period of time leads to muscle fatigue comprising a central and peripheral component, which demands a gradually increasing effort. However, the brain contribution to the enhancement of effort to cope with progressing fatigue lacks a complete understanding. The intermittent motor tasks (IMTs) closely resemble many activities of daily living (ADL), thus remaining physiologically relevant to study fatigue. The scope of this study is therefore to investigate the EEG-based brain activation patterns in healthy subjects performing IMT until self-perceived exhaustion. Fourteen participants (median age 51.5 years; age range 26-72 years; 5 males) repeated elbow flexion contractions at 40% maximum voluntary contraction by following visual cues displayed on an oscilloscope screen until subjective exhaustion. Each contraction lasted for approximately 5 s with a 2-s rest between trials. The force, EEG, and surface EMG (from elbow joint muscles) data were simultaneously collected. After preprocessing, we selected a subset of trials at the beginning, middle, and end of the study session representing brain activities germane to mild, moderate, and severe fatigue conditions, respectively, to compare and contrast the changes in the EEG time-frequency (TF) characteristics across the conditions. The outcome of channel- and source-level TF analyses reveals that the theta, alpha, and beta power spectral densities vary in proportion to fatigue levels in cortical motor areas. We observed a statistically significant change in the band-specific spectral power in relation to the graded fatigue from both the steady- and post-contraction EEG data. The findings would enhance our understanding on the etiology and physiology of voluntary motor-action-related fatigue and provide pointers to counteract the perception of muscle weakness and lack of motor endurance associated with ADL. The study outcome would help rationalize why certain patients experience exacerbated fatigue while carrying out mundane tasks, evaluate how clinical conditions such as neurological disorders and cancer treatment alter neural mechanisms underlying fatigue in future studies, and develop therapeutic strategies for restoring the patients’ ability to participate in ADL by mitigating the central and muscle fatigue.

## 1 INTRODUCTION

Fatigue interferes with task performance in multiple ways—slowing down, improper execution, or failure to accomplish. Furthermore, unpleasant sensations such as pain, discomfort, and increased effort can accompany the muscle fatigue. In healthy population, the consequences due to fatigue may be reflected as reduced physical performance, difficulty to carry out strenuous or prolonged activity, and demand for extra effort to perform everyday tasks, which will subside after a period of rest or decreased activity. Nevertheless, in individuals suffering from muscular, neurological, cardiovascular, and respiratory diseases and those who are aging or frail, the effects of fatigue are amplified and posing restrictions on daily life (Taylor et al., 2016).

Physical fatigue can be viewed as the decline in the voluntary-force-generating capacity of the neuromuscular system induced by a physical activity (Fry et al., 2017). It stems from active skeletal muscles that are involved in the peripheral processes as well as supraspinal mechanisms within the brain, thus comprising physiological and psychological aspects (Berchicci et al., 2013; Gandevia, 2001). The fatigue processes at or distal to the neuromuscular junction is referred to as peripheral fatigue and those attributed to the central nervous system affecting the neural drive to the muscle as central fatigue (Bigland-Ritchie et al., 1978).

The force generated by muscle contractions depends on the motor unit (MU) firing in the neuromuscular pathway: the cessation or slowing down of MU firing marks fatigue, whereas the increased recruitment of MUs serves as the compensatory mechanism. The central fatigue changes the MU firing by altering the intrinsic properties of motoneurons, feedback from the sensory input, and descending drive, which in turn modifies the strength and timing of muscle contractions. Besides, a sensation of discomfort and fatigue is facilitated by the firing of group III/IV muscle afferents. Furthermore, the modulation of brain neurotransmitters impacts the task endurance performance (Taylor et al., 2016). Therefore, the neural contribution to muscle fatigue is multi-pronged as depicted in Fig. 1 and gaining better insights into this mechanism holds paramount importance. Even though the physical fatigue is influenced by the supraspinal mechanisms within the brain, our understanding on how fatigue modulates the brain activity is far from complete.

**Figure 1.**
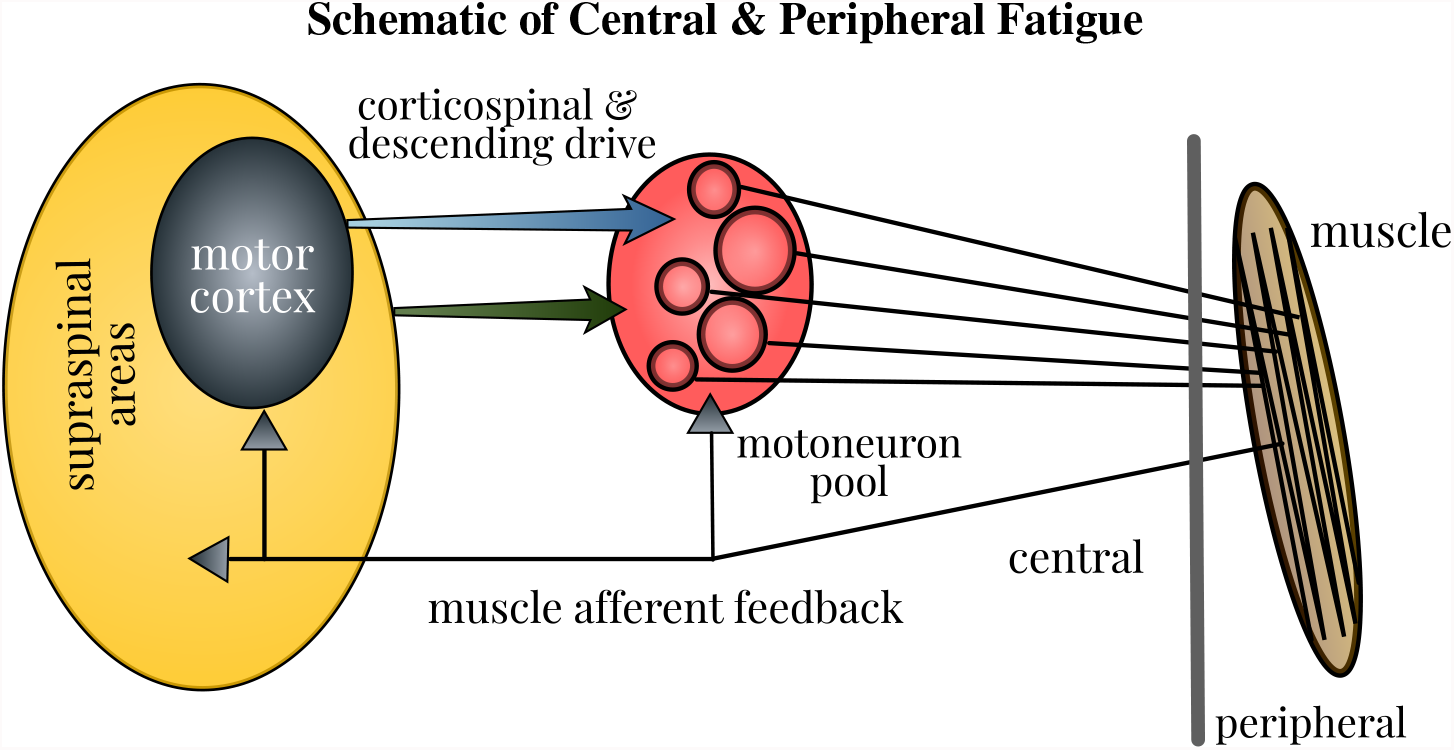
Schematic of neural contributions to muscle fatigue in single-joint contractions. Motoneuron firing is affected by the properties of motoneurons, muscle afferent feedback, and the corticospinal and other descending drive. Fatigue induces changes at all these levels, thus influencing the strength and timing of muscle contractions. Accordingly, the force generated by contraction of muscle fibers in the neuromuscular pathway is modified. Figure courtesy of (Taylor et al., 2016).

Aside from electrophysiological markers, fatigue can also be subjectively quantified by the perception of effort during physical activities, e.g., psychophysiological scale developed in (Borg, 1982). To understand the central and peripheral mechanisms underlying neuromuscular fatigue while performing isometric muscle contractions, several studies (Falvo et al., 2011; Pincivero, 2011; De Morree et al., 2012) have measured the perception of effort with the Borg category ratio scale.

The functional magnetic resonance imaging (fMRI) studies were dedicated to observe the changes in the cortical activation due to single-limb contractions until self-perceived exhaustion. The sustained maximal (Post et al., 2009; Steens et al., 2012) and submaximal contractions (Van Duinen et al., 2007; Benwell et al., 2007) with hand muscles progressively enhanced the activation in the motor cortex, supplementary motor area (SMA), and sensorimotor cortex. On the contrary, the repetitive fatigue task (Liu et al., 2005b) distinctly differs from the continuous one (Yang et al., 2010) in the sense that the former does not alter the blood-oxygen-level-dependent (BOLD) signal from the motor function cortices. The study findings would thus resolve an important research dilemma: do the motor-related cortices handle the fatigue induced by the repetitive and continuous task differently? Our fatigue study involving repetitive submaximal elbow flexion attempts to clarify whether the band-specific power of the electroencephalographic (EEG) signals recorded over specific cortical regions could be modulated by fatigue. To this end, we analyze the event-related spectral perturbation (ERSP), which is the estimation of post-onset and post-offset changes in the EEG spectrum with respect to the baseline (rest). The onset and offset refer to the time instant marking the application of a task-related stimulus and the cessation of the task, respectively. A more formal definition of ERSP is deferred to Section 4.

A few magnetoencephalographic (MEG) studies reported the effect of fatigue on the spectral power of signals recorded from the sensorimotor cortex. The post-movement beta rebound (PMBR) was reported to increase because of submaximal contractions until fatigue (Fry et al., 2017). Likewise, the muscle fatigue enhanced the beta- and gamma-band power of MEG signals while a motor task was carried out (Tecchio et al., 2006). Nevertheless, a repetitive maximal contraction fatigue task reduced the MEG alpha-band power from the sensorimotor and prefrontal areas (Tanaka et al., 2015). These contradictory results led us to speculate that even among the repetitive tasks, the influence of fatigue on the neural mechanisms may differ between the submaximal and maximal contractions. Our study is therefore intended to verify if the EEG spectral power changes due to fatigue caused by repetitive submaximal contractions agree with the MEG study findings.

There are ample evidences to believe that the fatigue task involving submaximal isometric contractions causes a significant rise in movement-related cortical potentials (MRCP) in brain regions that include primary motor cortex (M1), SMA, and premotor cortex, e.g., studies related to upper limb (Johnston et al., 2001; Guo et al., 2014; De Morree et al., 2012; Schillings et al., 2006) and lower limb (Berchicci et al., 2013). Similarly, a significant increase in the alpha- or beta-band EEG power in SMA, frontal, parietal lobe, and Brodmann area 11 was attributed to fatigue caused by cycling (Schneider et al., 2009; Hilty et al., 2011; Enders et al., 2016). By contrast, intermittent maximal and sustained submaximal contractions significantly reduced the alpha- and beta-band power measured at the EEG electrodes over the central and parietal lobes (Liu et al., 2005a; Nishihira et al., 1995). Thus, the findings related to band-specific EEG power changes due to a fatigue task remain inconclusive, which warrant further investigation.

Fatigue gradually sets in soon after the commencement of contractions, regardless of the fact that the individual continues the task execution (Barry and Enoka, 2007). It explains the reason why fatigue, task failure, and exhaustion are distinguishable (Berchicci et al., 2013). On this premise, we hypothesize that as the execution of repetitive submaximal contractions proceeds in time, the individual would experience increased levels of fatigue, and hence the modulation in the activation of motor-related cortices marking the fatigue level would also scale proportionally. Our study outcome would add knowledge of central mechanism of muscle fatigue as the experiment has been designed bearing the following objectives in mind. (i) The primary goal is to investigate how the EEG spectral power is altered with regard to graded fatigue levels, which has not been explored by past studies. (ii) The secondary goal is to help reach a consensus with the contradictory findings from fMRI, MEG, and EEG fatigue studies involving voluntary muscle contractions. (iii) The future goals are to make use of this knowledge to understand how the spectral power changes with fatigue are modified by a pathology and to design effective rehabilitation techniques targeting patients who have impaired movement functions.

## 2 METHODS

The current analysis was carried out on a previously collected dataset for which only the analysis on the muscle and force data was published (Cai et al., 2014).

### 2.1 Subjects

Fourteen healthy volunteers, of whom six were males, were enrolled in a study, where they were asked to perform an intermittent submaximal motor task repeatedly until they experienced severe fatigue to a point that they could not proceed any longer (Cai et al., 2014). The median age of the participants was 51.5 years and their age range was 26−72 years. Prior to the recruitment, the subjects were questioned whether they were depressed or not, and those who replied in the negative were only included in the study. The subjects were recruited through local advertisement. The approval for the study procedures was sought from the local Institutional Review Board and the subjects were required to provide a written informed consent. The demographics of the healthy cohort are presented in Table S1 (Supplementary Material).

### 2.2 Experimental Procedure

The participants first familiarized themselves with the submaximal elbow flexion task by attempting two isometric contractions with ≈ 30 −40% perceived maximal effort that lasted for a few seconds. After the warm-up activities, the Maximum Voluntary Contraction (MVC) force of isometric elbow flexion was recorded. Next, they performed the intermittent motor task (IMT) comprising repetitive elbow flexion at 40% MVC until they were subjectively exhausted.

#### 2.2.1 MVC Force

The subjects were seated with their forearm in a neutral position making an elbow joint angle of ≈ 100°. They were instructed to exert the maximal strength while performing two elbow flexion contractions using their dominant arm. The MVC force displayed on an oscilloscope screen was recorded with a data acquisition system (1401 Plus, Cambridge Electronic Design, Ltd., Cambridge, UK) in a computer. Simultaneously, the MVC force was also measured by a force transducer (JR3 Universal Force-Moment Sensor System, Woodland, CA). Out of the two MVC measurements, the highest value was selected for the subsequent analysis. If the two measurements differed by more than 5%, the MVC task had to be repeated to ascertain that the maximum effort was exerted during the contraction.

#### 2.2.2. Intermittent Motor Fatigue Task

The IMT was intended to contract the elbow flexor muscles until the subjects perceived severe fatigue. Each submaximal intermittent contraction set at 40% MVC was maintained for 5 s. The effort level to perform submaximal contractions was chosen to be 40% MVC to ensure that severe fatigue would be induced within 30 min leading to task failure (Cai et al., 2014). Note that this intensity of effort would be similar to mild-moderate day-to-day tasks (such as holding an infant, picking up groceries, etc.). The subjects were guided by visual cues associated with the initiation and cessation of contractions via an oscilloscope display in order to perform a series of contractions interleaved by a rest period of 2 s. The contractions were repeated until the individuals encountered self-perceived exhaustion.

#### 2.2.3. EEG Data Acquisition

Scalp EEG signals were recorded with a high-density 128-channel EEG data acquisition system (Electrical Geodesics, Inc. Eugene, OR, USA), in which the electrodes were arranged in a hat-like net and interconnected using nylon strings. Out of 128 channels, three were dedicated for EMG measurements from the muscles, namely *Biceps Brachii, Brachioradialis* (two elbow flexors), and *Triceps Brachii* (elbow extensor). A fourth channel was used for recording the elbow flexion force signal sensed by the force transducer. After soaking the electrode net in the electrolyte—one liter of distilled water added with 1.5 teaspoons of potassium chloride—it was mounted on the subject’s head. The connection quality of the electrodes was ensured by bringing their impedance values below 10 kΩ. The signal from each channel was amplified by a factor of 75000, band-pass filtered (0.01 Hz to 100 Hz), and digitized at a sampling rate of 250 Hz using NeuroScan system (Compumedic NeuroScan, Charlotte, NC) (Yang, 2008).

For more specific details on the experimental protocol, interested readers may refer to (Cai et al., 2014; Yang, 2008).

## 3 DATA PREPROCESSING PIPELINE

The 124-channel EEG data were preprocessed as outlined in Fig. S1 (Supplementary Material) with the standard tools available in EEGLAB 2021.0 software (Delorme and Makeig, 2004) prior to carrying out the ERSP analyses.

### Filtering and Noise Removal

The EEG signals were first band-pass filtered using a Hamming windowed sinc finite impulse response filter with the lower and higher cut-off frequencies of 1 Hz and 100 Hz, respectively. Next, they were filtered using a notch filter to remove the 60 Hz power-line noise and its higher harmonics. The noisy bursts in the filtered EEG data were corrected with Artifact Subspace Reconstruction (ASR) approach, which is an EEGLAB plug-in (Mullen et al., 2015). The threshold for burst detection criterion (specified as *k* in the ASR algorithm) was set as 20 as recommended in (Chang et al., 2019) based on a rigorous evaluation. Afterwards, the EEG signals were re-referenced to the common average, i.e., the arithmetic average of all electrodes (Nunez et al., 2006), which is advocated for EEG source estimation.

### Epoching and Data Cleaning

The continuous EEG data were then epoched as illustrated in Fig. 2 (left) with respect to the event-markers denoting the onset and offset of the elbow flexion task as follows. For analyzing the EEG pertaining to the steady contraction, the epoch limits were selected as 1.5 s before and 5 s after the task onset. To study the post-contraction EEG, the epoch limits were 4 s before and 2 s after the task offset. In the case of post-contraction, the baseline EEG (from 1.5 s before the task initiation up to the onset) from every epoch was used to replace the data within the time range, 4 s to 2.5 s, before the task offset in the respective epoch.

**Figure 2.**
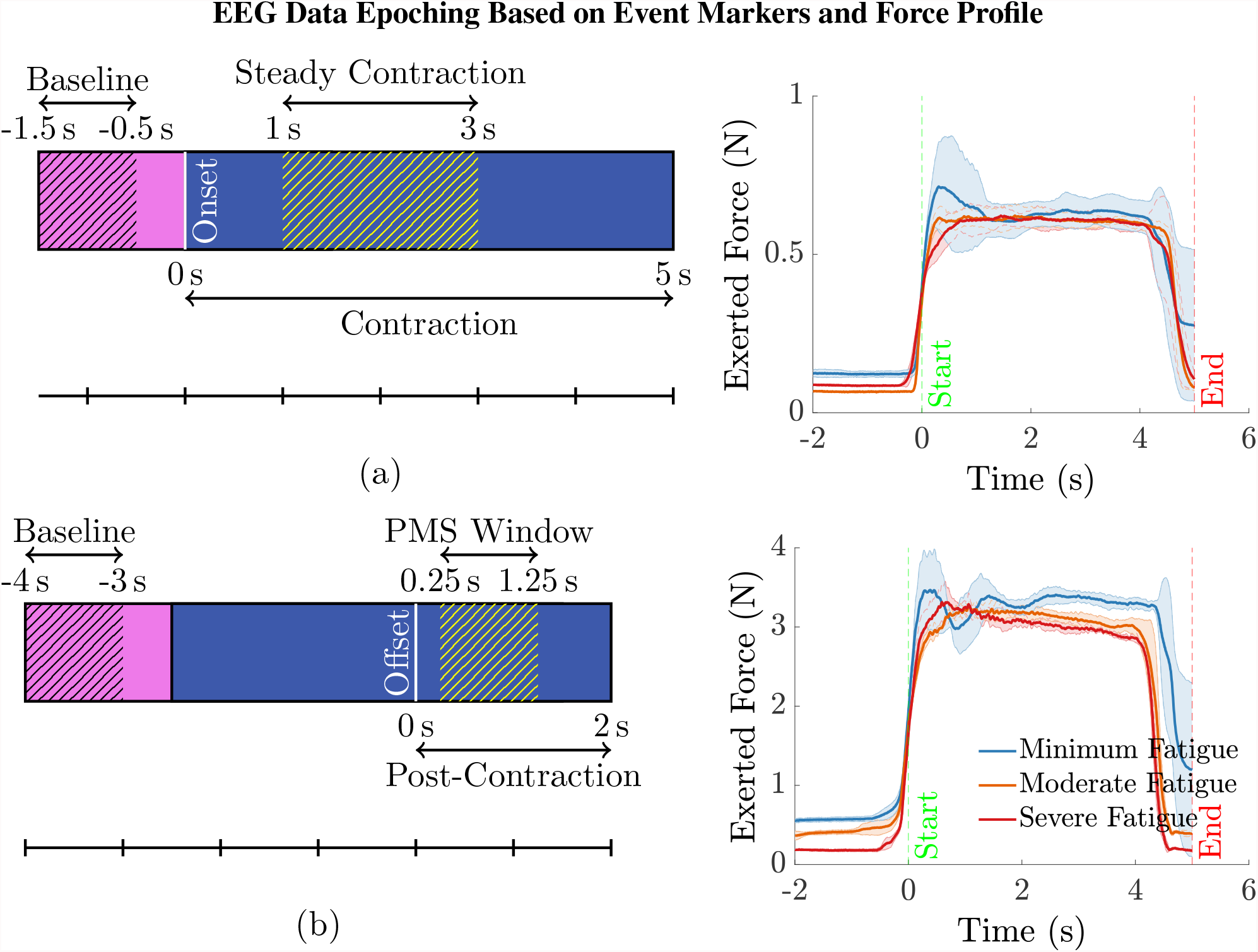
(a) To compute the ERSP for the steady contraction during 1 s to 3 s, EEG epochs were generated within the interval −1.5 s to 5 s, where 0 s represents the time instant of task initiation. The baseline EEG acquired during the period −1.5 s to −0.5 s was only used for the ERSP computation to preclude the data associated with movement anticipation. Similarly, the data within the interval 0 s to 1 s was disregarded as the subjects were ramping up to attain the target force of 40% MVC. (b) To study the ERSP for the time window of post-movement synchronization (PMS), i.e., 0.25 s to 1.25 s with 0 s denoting the time of task cessation, the data within the range −4 s to 2 s was 1epoched. Note that the data within the interval −4 s to −2.5 s was replaced by the respective baseline data in every epoch. Yet, the baseline data within the time span of −4 s to −3 s was only considered for calculating the ERSP for the aforementioned reason. The data recorded during the period 0 s to 0.25 s was discarded because the subjects required a brief duration to ramp down the force and reach the resting state. (right panel) The mean (solid line) and the standard deviation (shaded region) of the force recorded by a transducer during the intermittent elbow flexion trials performed by a male (bottom) and a female (top) representative healthy volunteer. For every individual, the total trials were divided into three equal sets, with the first, second, and the third set of trials representing the minimum, moderate, and severe fatigue condition. The task onset and offset are denoted by the event-markers, “ Start” and “ End” at 0 s and 5 s, respectively. Notice that almost all the subjects could maintain a steady contraction within the interval 1 s to 3 s after ramping up the force for about 1 s after the task onset.

The epoched data was visually inspected for the presence of noisy data. The epochs contaminated with noise, which would possibly affect the outcome of ERSP analysis, and those containing stim pulses (generated by supra-maximal stimulation of the muscle belly or motor nerve connecting the muscle to assess the peripheral fatigue, which is not investigated in this work) were removed.

### ICA Decomposition and Dipole Estimation

The multichannel EEG data free of noisy epochs and stim pulses was decomposed by an Independent Component Analysis (ICA) algorithm, namely Extended Infomax (Lee et al., 2000) (“ runica” in the EEGLAB toolbox), to estimate the underlying temporally independent EEG sources. The projection of EEG source signals to the scalp surface gave rise to scalp maps. We employed an EEGLAB plug-in, namely DIPFIT (Oostendorp and Van Oosterom, 1989), to identify the location of equivalent dipoles that could best describe the scalp maps. For this purpose, the boundary element spherical head model from the standard Brain Electrical Source Analysis [BESA (BESA GmbH, Gräfelfing, GE)] was adopted. Since the subject-specific digitized electrode positions were not available, the standard Montreal Neurological Institute (MNI) space that contains the dipole locations was applied to the head model.

### Artifactual IC Removal

The degree of resemblance between an independent component (IC) scalp map and the projection of the equivalent dipole is quantified using the residual scalp map variance (RV); for instance, the best-fitting model has the smallest RV. The ICs were classified with an artificial neural network framework into two categories: (i) “ non-brain-related ICs” representing electrical artifacts stemming from muscle, eye, heart, line noise, and channel noise and (ii) “ brain ICs” accounting predominantly for activities originating within the brain. The classifier was implemented with an EEGLAB plug-in (Pion-Tonachini et al., 2019), known as ICLabel. The ICs designated as non-brain-related and those having RV values exceeding 20% were discarded. This process called IC pruning will help remove artifactual IC components that are spatially stereotypical. With the retained neural ICs (RV*≤* 20%), the EEG signals were reconstructed to produce ICA-cleaned EEG for further analysis.

## 4. DATA ANALYSIS

We conducted the following ERSP analyses. (i) **Channel-Level ERSP Analysis**. It was performed directly with the EEG data recorded using electrodes mounted on the scalp over brain regions of interest. (ii) **Source-Level ERSP Analysis**. First, the ICs were estimated by decomposing the scalp EEG data with the Extended Infomax ICA algorithm (Bell and Sejnowski, 1995). Next, the ICs were clustered with the *k*-means algorithm from the MATLAB Statistics Toolbox based on the location and orientation of the equivalent dipoles to obtain physiologically and anatomically separate brain sources. The parameters of *k*-means were selected as the default values, i.e., the number of clusters and the standard deviation to decide the outliers were eight a<u>nd th</u>ree, respectively. The rule of thumb to determine the number of *k*-means clusters is suggested as 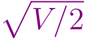 in p. 365, Mardia et al. (1980), where *V* is the number of data points to be clustered. In our case, this value coincides with the default choice in EEGLAB: on average there will be one IC component per subject per cluster. Finally, the analysis was carried out with the plausible EEG sources (IC clusters) located within motor-related cortical regions. In both cases, we performed the ERSP analysis with the EEG data corresponding to the steady elbow flexion and post-contraction [hatched blue region in Fig. 2 (a) and (b), respectively]. By visually inspecting the force data in Fig. 2 (right) that was simultaneously measured with the force transducer, the steady-contraction period was determined to be between 1 s and 3 s after the task initiation. The EEG data within the interval 0 s to 1 s [unhatched blue region in Fig. 2 (a)] was excluded from the analysis since the subjects were ramping up to reach certain target force that was set for the IMT trials. Likewise, while performing the ERSP analysis for post-contraction, the data recorded from 0 s to 0.25 s after the task offset [unhatched blue region in Fig. 2 (b)] was disregarded because it would contain remnant task-related EEG as the subjects were ramping down to rest after cessation of the task. The EEG acquired during the time interval immediately before the task onset, i.e., −0.5 s to 0 s [unhatched pink region in Fig. 2 (a)], was not considered as the baseline data, because this data segment would be associated with movement anticipation and hence does not represent the ideal state of rest. For the same rationale, the baseline data from −3 s to −2.5 s with respect to the task offset [unhatched pink region in Fig. 2 (b)] was not included in the ERSP calculation.

We computed the ERSP based on the formal definition provided in (Fuentemilla et al., 2006; Delorme et al., 2007; Meltzer et al., 2008) for the EEG data in a frequency band of interest during the steady contraction or post-contraction to compare the power spectral density (PSD) between the fatigue conditions at the channel or source level. Furthermore, we implemented the framework illustrated in Fig. S2 (Supplementary Material) to make statistical inferences. Recall from Section 2.2.2 that the healthy volunteers repeated the elbow flexion until they experienced severe fatigue (SevFatg), which eventually led to the cessation of intermittent contractions. We suppose that there would have been a steady progression of fatigue as the subjects prolonged the IMT execution and ultimately they reached SevFatg just before discontinuing the motor task. On this premise, we divided the entire EEG data into three sets with equal number of trials by preserving the order to maintain the correspondence to minimum fatigue (MinFatg), moderate fatigue (ModFatg), and SevFatg. For the ERSP analysis, we selected only the first, middle, and the last 10% of trials from the first, second, and third set of trials, respectively, to ensure that the data germane to the three fatigue conditions are distinctly disparate. In our EEG dataset, 10% of trials from every fatigue condition varies between the subjects within the range [5, 9] and has a median value of ≈ 8.

### Statistical Framework

For notational convenience in the sequel, the subject (or component) index *j* and the fatigue condition are omitted in (1)–(4). The mean event-related spectrum (ERS^1^) is the average data power spectrum across *M* trials computed for the sliding time windows centered at *t* in each trial, given by

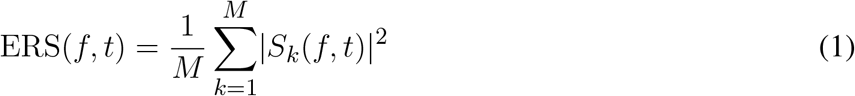

with *S*_*k*_(*f, t*) being the spectral estimate for trial *k* at frequency *f* and time instant *t*. In the present setting, *M* denotes the total number of trials for a given subject (channel-level) or component in an IC cluster (source-level) corresponding to any one of the three fatigue conditions. The term | *S*_*k*_(*f, t*) |^2^ represents the power spectrum for the *k*-th trial of subject (or component) *j*. We apply the gain model, which is the default choice in EEGLAB software that divides the contraction-induced spectral activity by the pre-contraction spectral activity. For a given frequency band, the absolute ERSP is computed by dividing the ERS power in (1) computed for the steady contraction or post-contraction at each time-frequency point by the average spectral power in the pre-contraction baseline period at the same frequency.

As pointed out in (Grandchamp and Delorme, 2011), the log-transformed measure of ERSP is endowed with the following merits: (i) The EEG signal distribution, in general, is skewed. By taking its logarithm, the distribution is made more normal, and hence it becomes suitable for parametric inference testing. (ii) The log-transformed power values help visualize a wider range of power variations, whereas its absolute value leads to the masking of power changes at high frequencies by those at low frequencies. Therefore, the ERSP analyses and the results reported here are based on the log-transformed ERSP in (2) and its time-averaged quantity in (3), both expressed in decibels (dBs):

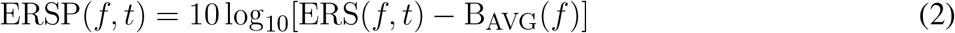

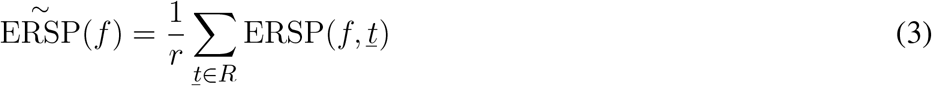

where

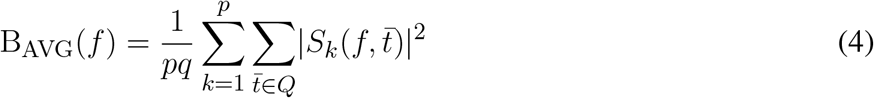

and *r* denotes the cardinality of the set *R* of steady-or post-contraction time points. In (4), B_AVG_(*f*) denotes the pre-contraction spectral power as a function of *f* averaged over a total of *q* time points belonging to the set *Q* in the baseline period across *p* trials. We derived a common baseline for every subject (or component), i.e., the power spectra of the pre-contraction data for the *j*-th subject (or component) were averaged across the trials, time points, and fatigue conditions. The rationale for using a common baseline across fatigue conditions is to counteract the effect due to fatigue-induced changes in the baseline to the best possible extent. Consequently, *p* in (4) represents the sum of selected number of trials for a subject (or component) under MinFatg, ModFatg, and SevFatg. We vertically concatenated the row vectors 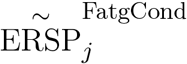 for *j* = 1, 2, …, *N* to construct a matrix of size *N* × *f* as given by

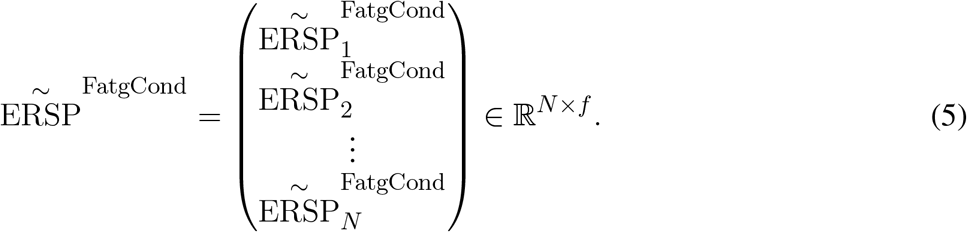

To draw statistical inferences, we conducted the surrogate permutation test with the following: (i) log-transformed ERSP matrices 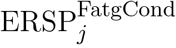 defined in (2) for frequencies in the interval 3 Hz to 90 Hz and time period shown in Fig. 2 (left); (ii) log-transformed time-averaged ERSP vectors that are vertically concatenated as in (5) to build the matrix 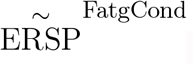. Subsequently, we applied a false discovery rate (FDR) correction test in both scenarios to assess how the FDR was controlled for multiple comparisons (Lage-Castellanos et al., 2010). Finally, we performed a one-way analysis of variance (ANOVA) test after averaging the PSD values in 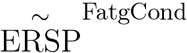 over the EEG frequency bands^2^ of interest, which is henceforth referred to as 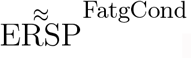

## 5 STATISTICAL INFERENCES

### Permutation Test & FDR Correction

We studied the log-transformed matrices 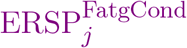 and the vertical concatenation of log-transformed time-averaged row vectors 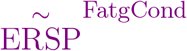 with the surrogate permutation test from the EEGLAB toolbox. We adopted the permutation test (Pesarin, 1992) that draws samples without replacement (Rimbert et al., 2019), because the EEG data may not conform to the normality assumption required by many statistical approaches. Moreover, the test is deemed suitable for our small sample size, i.e., 14, and in scenarios where the data contain outliers, which is more likely our case. We selected the following values for the permutation test: *p* < 0.05 and 2000 permutations. The test results validated the differences among the three fatigue conditions in terms of ERSP values at each time-frequency point. We applied the false discovery rate (FDR) correction to control the type I errors in null hypothesis testing when conducting multiple comparisons with a *p*-value threshold of 0.05. The reason for employing FDR is that it is less conservative compared to family-wise error rate (FWER) controlling procedures and have greater power (Benjamini and Hochberg, 1995).

### One-Way ANOVA & Tukey-Kramer Test

The one-way ANOVA (Howell, 2012) test was performed with 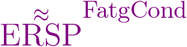 obtained by averaging the log-transformed ERSP over the steady-/post-contraction time interval and the frequency band of interest to verify whether their means are different. While ANOVA is considered to be robust when the normality assumption is violated (Kirk, 2012), we rely on the log transformation that enables the data distribution to be made normal, thus rendering it suitable for parametric inference testing. We conducted the multiple comparison test (post-hoc analysis), namely Tukey’s honestly significant difference procedure (Tukey, 1949), when the *p*-value for the ANOVA was statistically significant to explore the difference between the three group means while controlling for the FWER.

## 6 RESULTS

The ERSP analysis was carried out across the three fatigue conditions and the results were reported for the following four schemes: (i) channel-level ERSP for steady contraction; (ii) source-level ERSP for steady contraction; (iii) channel-level ERSP for post-contraction; and (iv) source-level ERSP for post-contraction.

### 6.1 Selected Channels & IC Clusters

Out of 124 EEG channels, we selected six channels, namely FC3, C1, C3, C5, CP1, and CP3, corresponding to the scalp electrodes mounted over the contralateral cortical regions associated with the IMT (elbow flexion) for the channel-level analysis. Note that all the 14 healthy participants were right-handed. To perform the source-level analysis, first, the ICs were estimated from the multichannel EEG signals by the ICA algorithm and fitted with the equivalent dipoles via DIPFIT toolbox. Second, the ICs across the subjects were clustered with *k*-means algorithm based on the location and orientation of equivalent dipoles as described in Section 3. Third, the scalp maps of cluster centroids and the clusters of equivalent dipoles were visually inspected to identify the plausible neural sources that were located in close proximity to the cortical regions of interest responsible for the upper extremity motor task. The dipole clusters of interest (blue dots) and the respective cluster centroids (red dots) were shown for the steady-contraction EEG in the first row and for the post-contraction EEG in the second row of Fig. 3.

**Figure 3.**
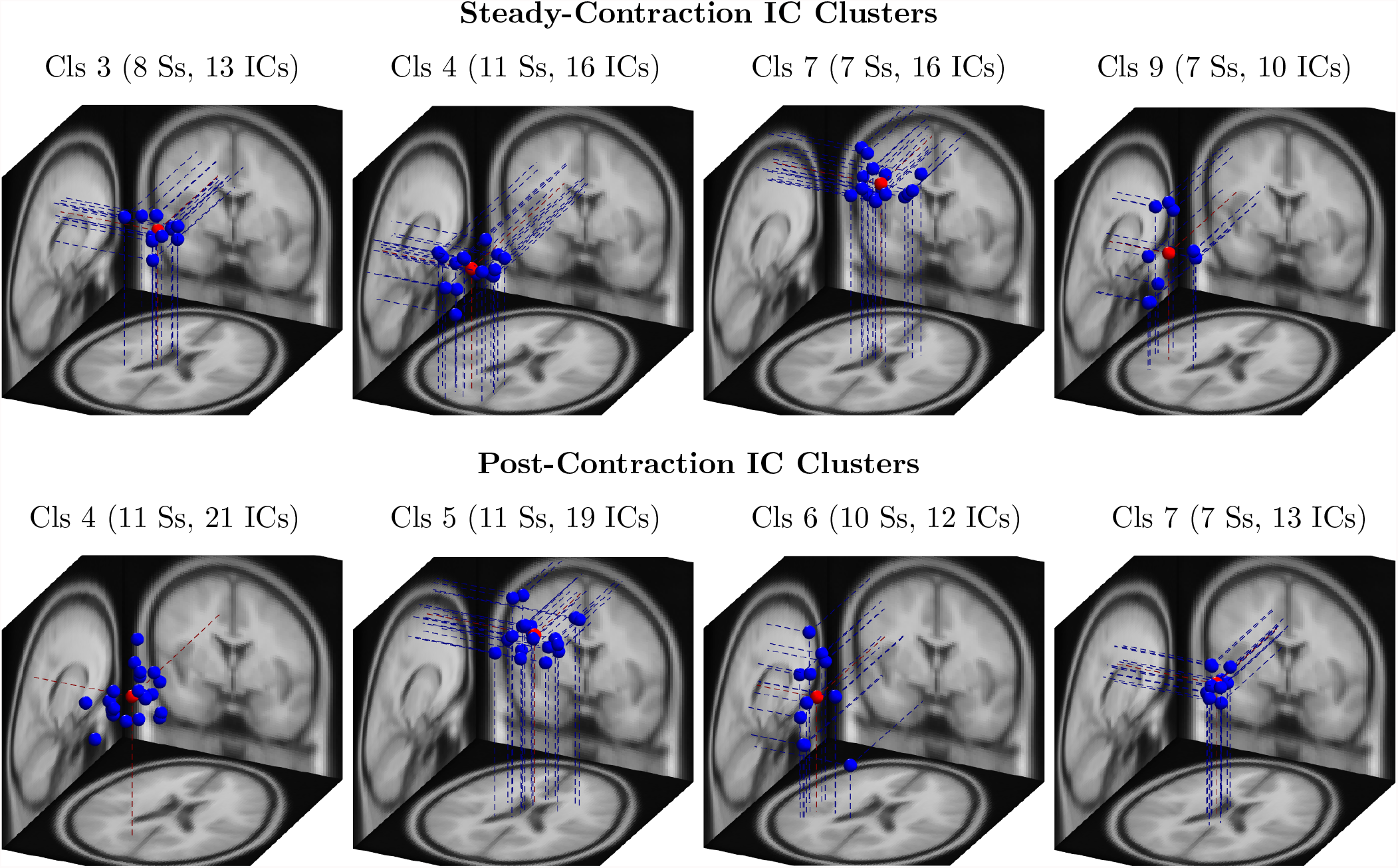
The clusters of equivalent dipoles were identified based on their location and orientation with the *k*-means algorithm as described in Section 4 for source-level analysis. Each IC cluster (Cls) is assigned an arbitrary index for identification; the number of ICs constituting the cluster and the number of associated subjects (Ss) are provided within the parentheses. The IC clusters represent plausible neural sources estimated from steady-contraction EEG (row 1) and post-contraction EEG (row 2), and they are located in the following brain regions: (row 1) Cls 3 - left cingulate gyrus; Cls 4 - left superior parietal gyrus Cls 7 - right paracentral lobule; Cls 9 - left transverse temporal gyrus; (row 2) Cls 4 - left precuneus; Cls 5 - right paracentral lobule; Cls 6 - left transverse temporal gyrus; and Cls 7 - left cingulate gyrus. The dipoles are marked with blue dots and the cluster centroids estimated by the *k*-means with red dots.

### 6.2 ERSP Represented as Time-Frequency Image

For a given fatigue condition, we computed the log-transformed ERSP measure in (2) from an EEG channel of interest for all the 14 subjects in the channel-level analysis. The EEG channel was replaced by an IC cluster of interest, while performing the source-level analysis with the ERSPs computed for the components of the cluster. We generated separate sets of 2-D patterns obtained by averaging 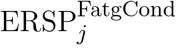 over *N* subjects (components), denoted as 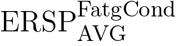 in the sequel, for the steady and post-contraction using both analyses. We reiterate that since our objective is to compare the PSDs across MinFatg, ModFatg, and SevFatg, a subject- or component-wise common baseline needs to be derived as in (4) for calculating the ERSP to minimize the effect of fatigue on the baseline. Among the six selected channels, namely FC3, C1, C3, C5, CP1, and CP3, representative ERSP patterns from CP1 and CP3 are displayed in Fig. 4 (row 1 & 2) for the steady contraction. Likewise, typical ERSP patterns from C3 and C5 are shown for the post-contraction in Fig. 5 (row 1 & 2). We performed the source-level analysis on the following four IC clusters^3^ because they were located within the cortical regions associated with the execution of motor tasks and are shown in Fig. 3: IC Cluster 3, 4, 7, and 9 for the steady-contraction data and IC Cluster 4, 5, 6, and 7 for the post-contraction data. A sample set of patterns from the source-level analysis is shown under each category—steady contraction in Fig. 4 (row 3) and post-contraction in Fig. 5 (row 3). The task (elbow flexion) onset and offset are marked in all the patterns with a dotted vertical line in Fig. 4 and Fig. 5, respectively. The last column in both Fig. 4 and 5 is the pictorial representation of the outcome of surrogate permutation test subject to the FDR correction with *p* < 0.05. Interestingly, in all instances from the channel- and source-level analysis for the steady or post-contraction, we observed statistically significant differences across MinFatg, ModFatg, and SevFatg at a large collection of time-frequency points within the theta-, alpha-, and beta-band frequency range.

**Figure 4.**
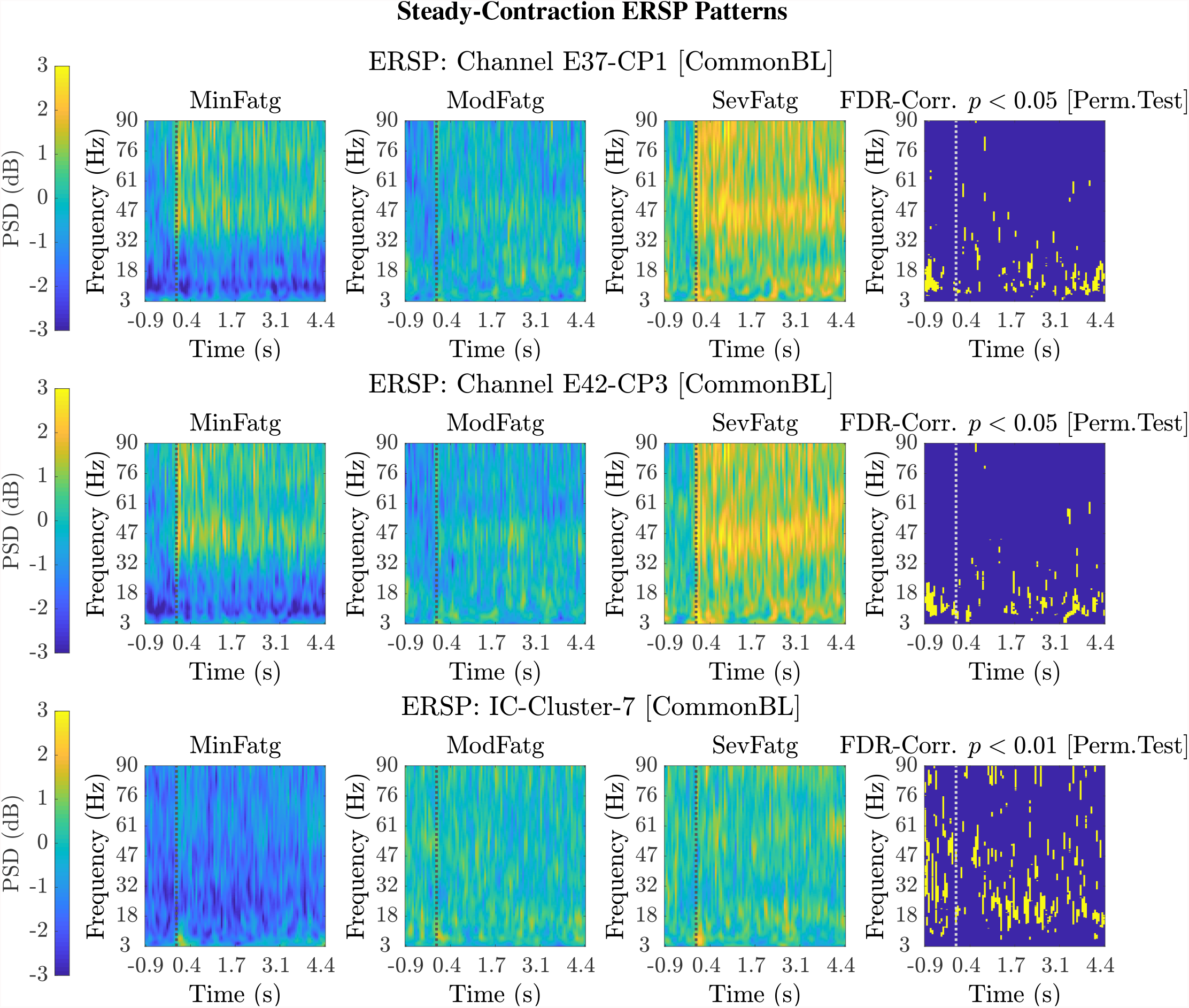
Representative 2-D patterns 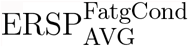 generated for the EEG data recorded from two scalp electrodes, CP1 and CP3, during the steady contraction are depicted in the top and middle row, respectively. For the source-level analysis, the ICA decomposition was performed on the multichannel EEG signals and the IC clusters were estimated. The ERSP patterns are shown in the bottom row for IC Cluster 7 that comprises ICs located near the right paracentral lobule. Both the channel-level and source-level ERSP analyses reveal that the PSD computed in the theta, alpha, and beta frequency band scales with the fatigue level. The results from the permutation test followed by the FDR correction with a threshold *p*-value of 0.05 (except for IC Cluster 7 with *p* < 0.01) are displayed in the last column in every row. The dotted vertical line in each ERSP pattern denotes the onset of the elbow flexion task.

**Figure 5.**
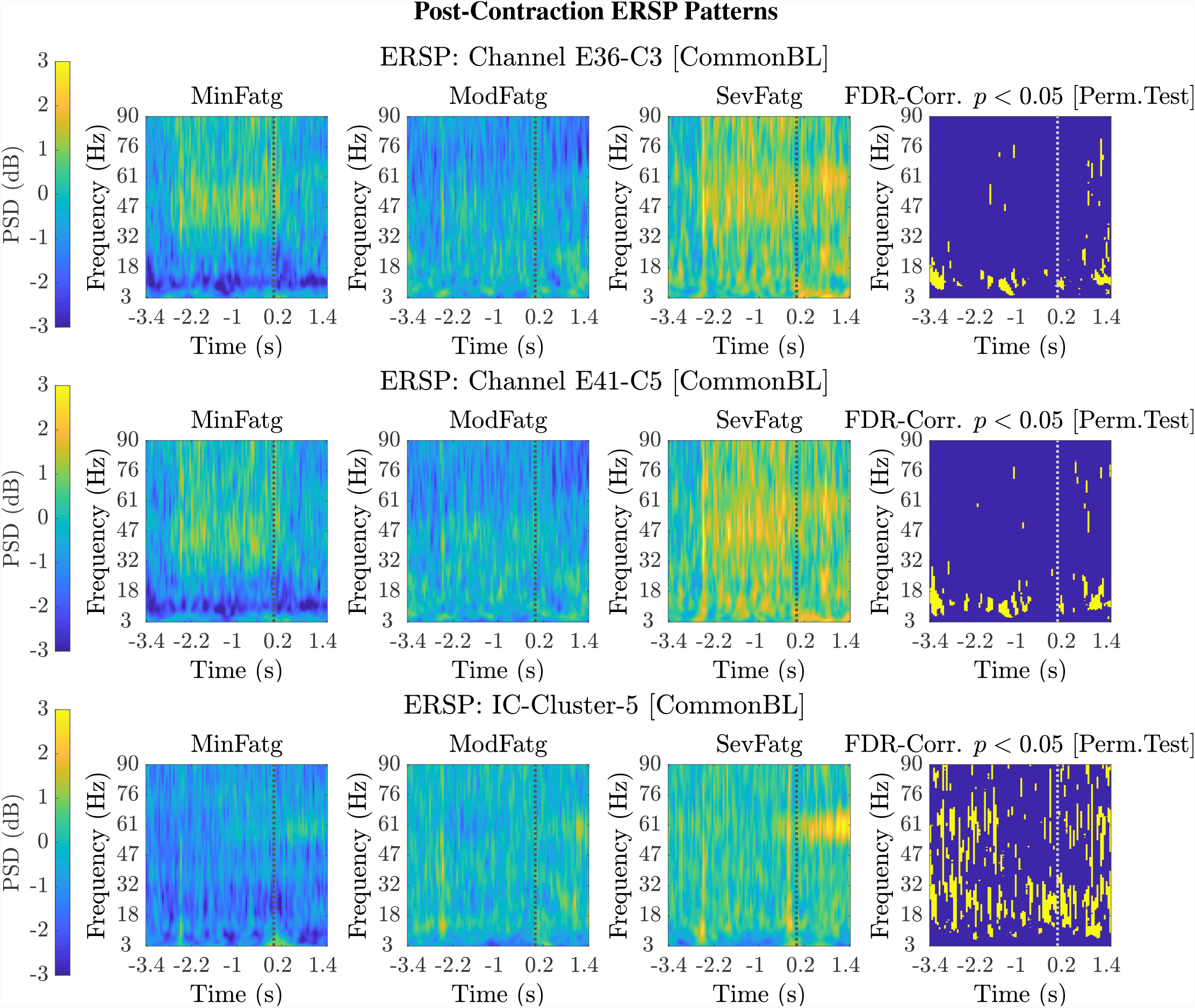
Typical 2-D patterns 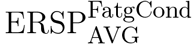 for the EEG signals recorded during post-contraction with the scalp electrodes, C3 and C5, are shown in the top and middle row, respectively. The set of ERSP patterns displayed in the bottom row corresponds to IC Cluster 5 situated in the right paracentral lobule. The ERSP patterns offer evidence for a monotonic increase in the theta-, alpha-, and beta-band PSD with fatigue. The FDR-corrected *p*-values less than 0.05 threshold from the permutation test are depicted in the last column to indicate the statistical significance. The dotted vertical line in each ERSP pattern represents the offset of the elbow flexion task.

### 6.3 ERSP Averaged Over Task-Related Time Intervals

The PSD values of time-frequency points in 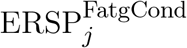 were averaged over time within the steady- (1 s to 3 s) or post-contraction (0.25 s to 1.25 s) interval to deduce 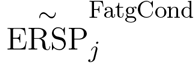 for *j*-th subject (component) to help investigate how the PSD modulates across the fatigue conditions in the steady- or post-contraction interval as the frequency varies within the range 3 Hz to 90 Hz. Simply put, 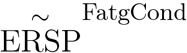 for a channel or neural source is the time-averaged PSD for different subjects (components) expressed as a function of frequency. For the steady contraction, the 1-D ERSPs are plotted for channel CP1 and CP3 in the left and right panel of Fig. 6, respectively. Fig. 7 (left panel) shows the plots for IC Cluster 7, which comprises ICs located within a motor-related cortical area (right paracentral lobule) as depicted in Fig. 7 (right panel). Notice that for CP1 and CP3, the PSD averaged over time increases with fatigue maintaining a statistical significance (*p* < 0.05) in the theta, alpha, and beta band per the permutation test with FDR correction for multiple comparisons. Also noteworthy is that in IC Cluster 7, even though the mean PSD pertaining to ModFatg and SevFatg seems to overlap at around 25 Hz, the permutation test with FDR correction could detect the difference at those frequencies. In like manner, the channel- and source-level 1-D ERSPs for post-contraction are exemplified in Fig. 8 and 9, which underscore our key finding, i.e., in both analyses, the time-averaged PSD increases as the perceived fatigue escalates in the theta, alpha, and beta band. For post-contraction, the plots were generated from 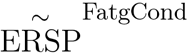 for channel C3 and C5 [Fig. 8 (left and right panel)] as well as IC Cluster 5 [Fig. 9 (left panel)] containing ICs positioned in a cortical region related to motor function (right paracentral lobule) [Fig. 9 (right panel)].

**Figure 6.**
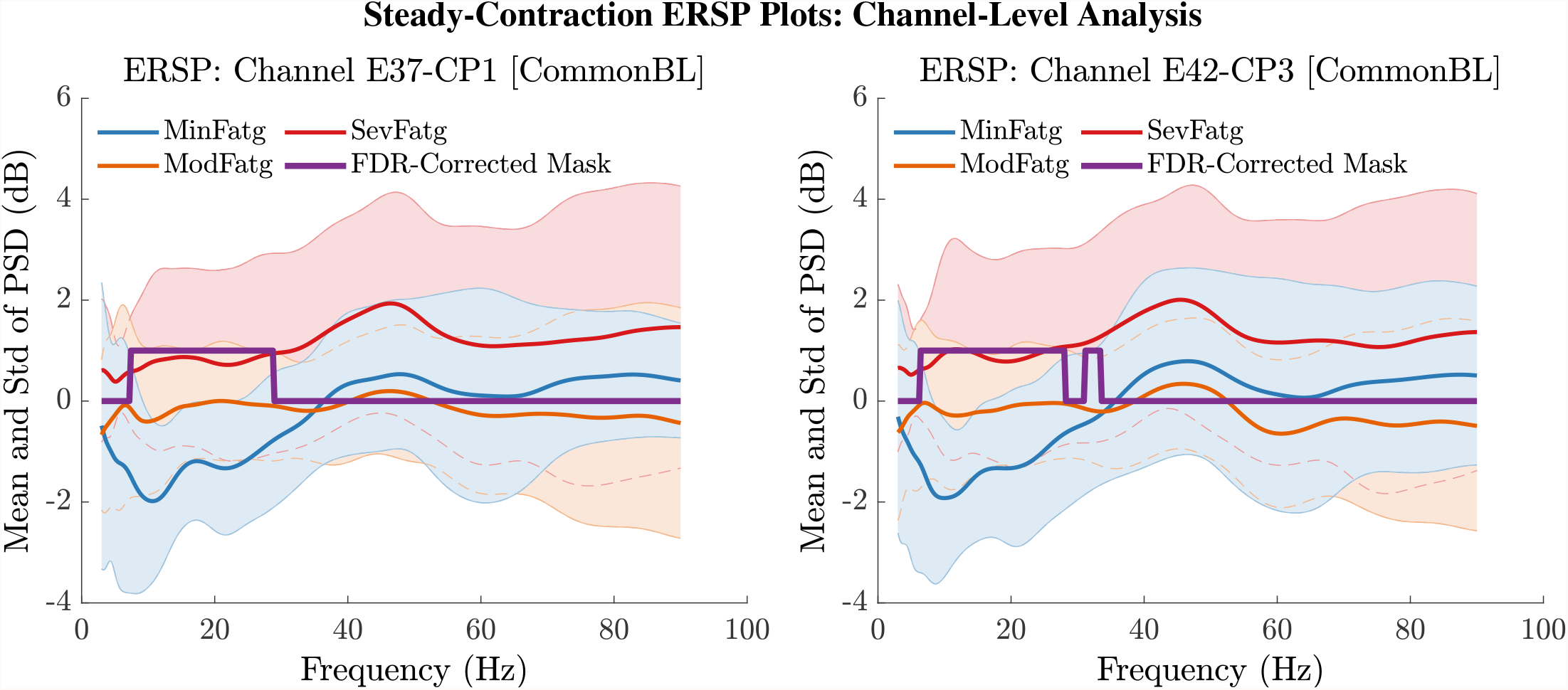
The ERSP values at time-frequency points were averaged over the steady-contraction period of 1 s to 3 s for EEG signals. The plots produced from 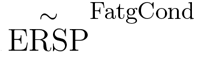 help understand how the time-averaged PSD computed for the scalp EEG varies with IMT-induced fatigue. The mean and standard deviation (std) of PSD pertaining to the three fatigue conditions are expressed as functions of frequency within the range 3 Hz to 90 Hz for the EEG channel CP1 (left panel) and CP3 (right panel). The frequencies at which the PSD differs across the fatigue conditions with a statistical significance (*p* < 0.05) are assigned a value one and the rest of the frequencies a value zero by the FDR-corrected mask.

**Figure 7.**
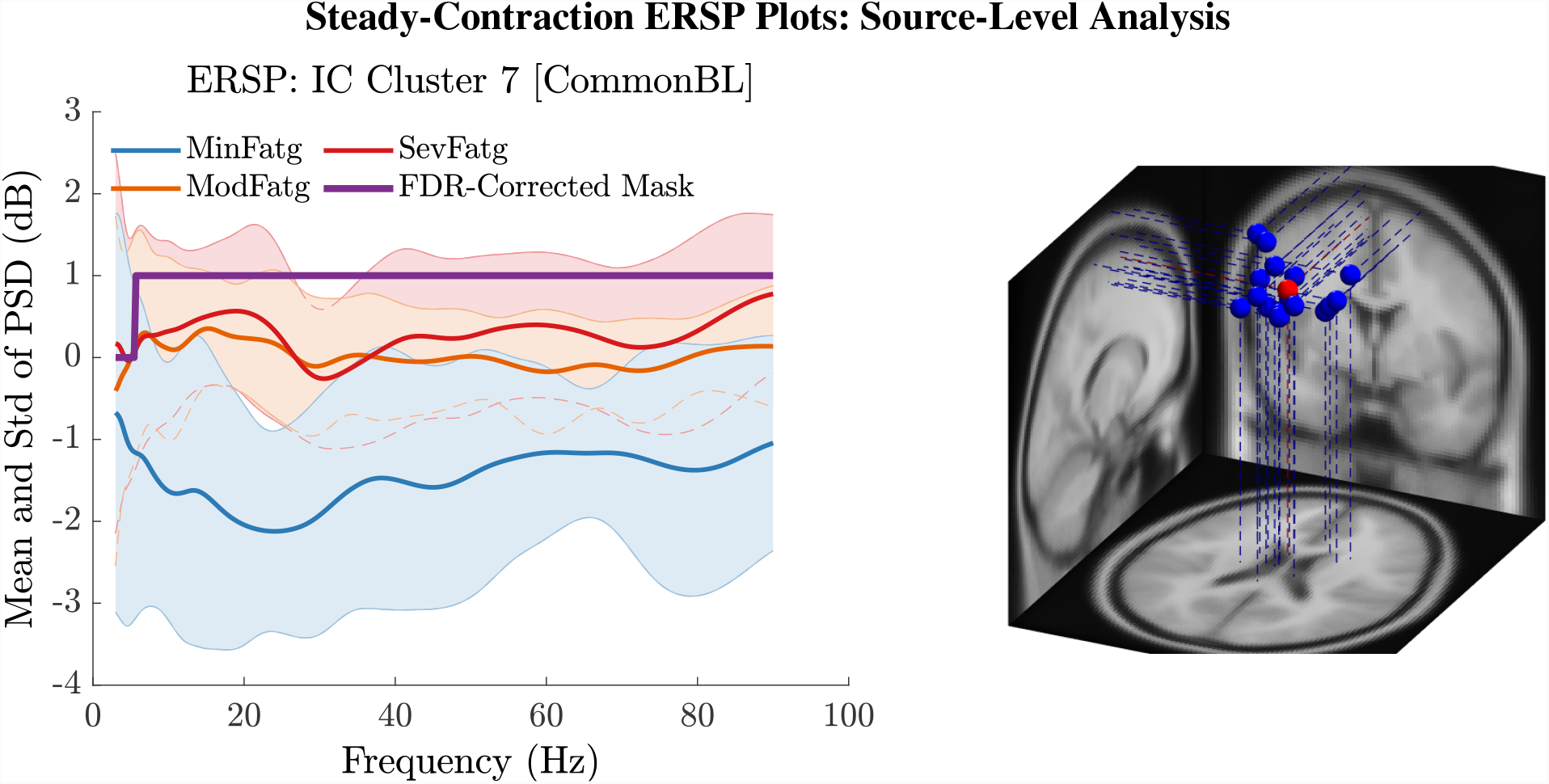
The steady-contraction ERSP averaged over time, 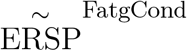, portrays how the PSD for IC Cluster 7 modulates across MinFatg, ModFatg, and SevFatg, as a function of frequency (left panel). The FDR-corrected mask indicates that a statistically significant (*p* < 0.05) difference in PSD between the three fatigue conditions is maintained through out the EEG frequency spectrum except the delta band. The brain sources (ICs) marked as equivalent dipoles (blue dots) in Cluster 7 and the cluster centroid (red dot) are located in vicinity to the right paracentral lobule (right panel).

**Figure 8.**
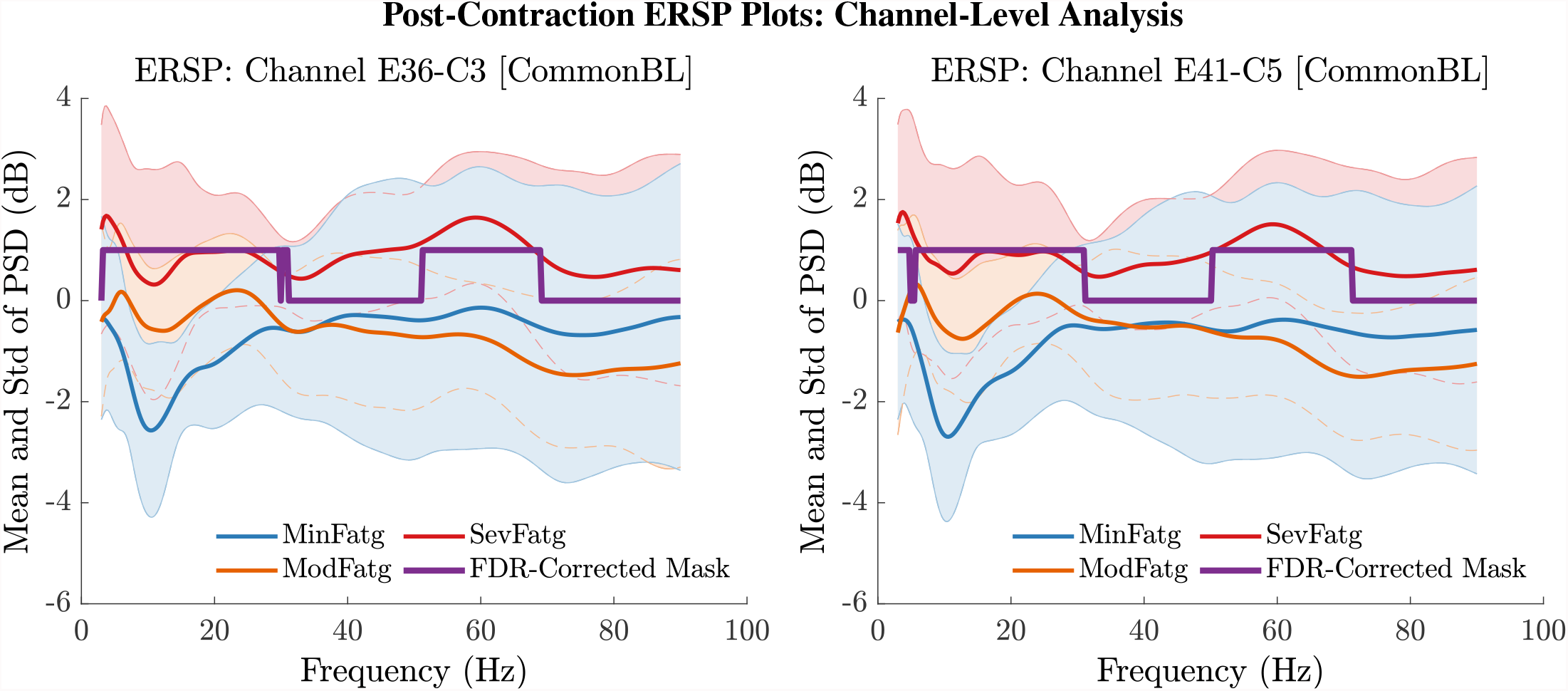
The PSD values in 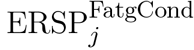 were averaged over the post-contraction interval of 0.25 s to 1.25 s to obtain the mean and std of time-averaged PSD for frequencies from 3 Hz to 90 Hz for the EEG channel C3 (left panel) and C5 (right panel). The FDR-corrected mask illustrates that the PSD values differ between the fatigue conditions at theta-, alpha-, and beta-band frequencies with a statistical significance (*p* < 0.05).

**Figure 9.**
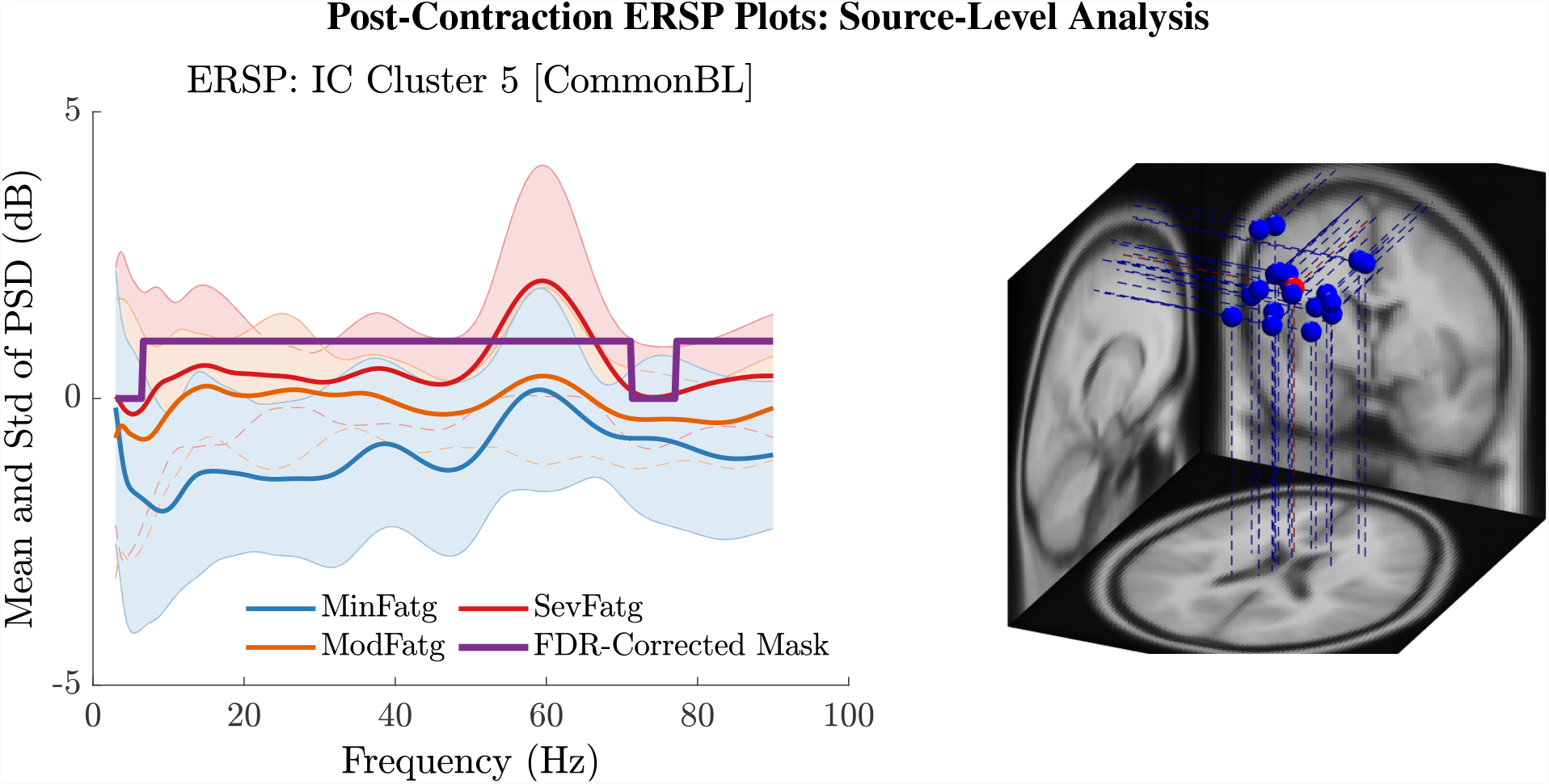
The post-contraction ERSP averaged over time, 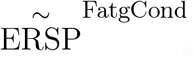, shows the PSD variations across the fatigue conditions for the frequency range 3 Hz to 90 Hz in IC Cluster 5 (left panel). The FDR-corrected mask indicates that the time-averaged ERSP differs between fatigue levels with a statistical significance (*p* < 0.05) in the entire frequency range except in the delta and high gamma band. The equivalent dipoles corresponding to ICs (blue dots) in Cluster 5 and its centroid (red dot) are seated within the right paracentral lobule or SMA (right panel).

### 6.4 ERSP Averaged Over Task-Related Time Intervals & Frequency Bands

We averaged 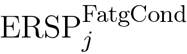 across the time and frequency points within the steady-/post-contraction period and a motor-related frequency band (theta, alpha, or beta), respectively. Thus, the band-specific PSD values corresponding to a time period were evaluated for MinFatg, ModFatg, and SevFatg, which are represented as a vector 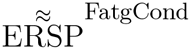 ERSP of length *N* for all the subjects (components). The statistical inferences help verify whether the averaged ERSP measure maintains a monotonic relationship with fatigue and differs between the three fatigue conditions with a statistical significance. To this end, we applied the one-way ANOVA to 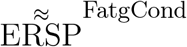 values estimated for the subjects (channel-level) or IC components (source-level). If the test results turned out to be overall significant (*p* < 0.05), then we conducted the post hoc Tukey’s test to determine the significance of differences between pairs of group means. For a given frequency band, the pairwise differences of time- and frequency-averaged PSD between the three fatigue groups were visually inspected with the box plots. For the steady contraction, the box plots are provided for two EEG channels—CP1 (row 1) and CP3 (row 2)—and IC Cluster 7 (row 3) in Fig. 10. Similarly, Fig. 11 contains the box plots of channel C3 (row 1), C5 (row 2), and IC Cluster 5 (row 3) for the post-contraction data. The first, second, and third column of Fig. 10 and 11 correspond to theta, alpha, and beta band, respectively. With a couple of exemptions (shaded in gray in Table 1),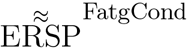 scales with fatigue for the EEG channels and IC clusters pertinent to the motor and sensory cortical areas during the steady- and post-contraction period.

**Table 1.** The one-way ANOVA test was conducted with the time-averaged theta-, alpha-, and beta-band PSD values 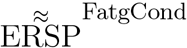 deduced from the scalp EEG and neural sources during the steady contraction. The *F* - and *p*-values of ANOVA and the *p*-values from the post-hoc analysis (if *p* < 0.05 from ANOVA) are listed. The light, medium, and dark shades of red denote that the PSD increases with fatigue with a statistical significance of *p* < 0.1, *p* < 0.05, and *p* < 0.001, respectively. The counter cases, where the PSD decreases as the fatigue increases without a statistical significance, are shaded in gray.

| Channel/<br>IC Cluster | Frequency<br>Band | ANOVA<br>$F$-Value | ANOVA<br>$p$-Value | Min-Mod Fatg<br>$p$-Value | Min-Sev Fatg<br>$p$-Value | Mod-Sev Fatg<br>$p$-Value |
| --- | --- | --- | --- | --- | --- | --- |
| FC3 | $\theta$ | 4.87e+00 | 1.30e-02 | 1.05e-01 | 1.12e-02 | 6.07e-01 |
| | $\alpha$ | 6.59e+00 | 3.43e-03 | 1.32e-01 | 2.33e-03 | 2.37e-01 |
| | $\beta$ | 5.73e+00 | 6.58e-03 | 4.17e-01 | 4.96e-03 | 1.08e-01 |
| C1 | $\theta$ | 4.02e+00 | 2.60e-02 | 3.50e-01 | 1.94e-02 | 3.35e-01 |
| | $\alpha$ | 8.30e+00 | 9.96e-04 | 8.79e-02 | 6.38e-04 | 1.54e-01 |
| | $\beta$ | 6.36e+00 | 4.06e-03 | 1.96e-01 | 2.75e-03 | 1.81e-01 |
| C3 | $\theta$ | 5.57e+00 | 7.48e-03 | 9.27e-02 | 6.00e-03 | 5.00e-01 |
| | $\alpha$ | 9.29e+00 | 5.02e-04 | 4.20e-02 | 3.30e-04 | 1.91e-01 |
| | $\beta$ | 4.86e+00 | 1.30e-02 | 3.22e-01 | 9.41e-03 | 2.34e-01 |
| C5 | $\theta$ | 6.29e+00 | 4.30e-03 | 6.78e-02 | 3.43e-03 | 4.67e-01 |
| | $\alpha$ | 6.94e+00 | 2.65e-03 | 1.70e-01 | 1.75e-03 | 1.57e-01 |
| | $\beta$ | 5.44e+00 | 8.25e-03 | 7.00e-01 | 8.08e-03 | 5.86e-02 |
| CP1 | $\theta$ | 3.66e+00 | 3.48e-02 | 2.43e-01 | 2.79e-02 | 5.49e-01 |
| | $\alpha$ | 1.04e+01 | 2.40e-04 | 3.04e-02 | 1.54e-04 | 1.55e-01 |
| | $\beta$ | 8.01e+00 | 1.22e-03 | 8.04e-02 | 8.00e-04 | 1.92e-01 |
| CP3 | $\theta$ | 5.02e+00 | 1.15e-02 | 1.65e-01 | 8.55e-03 | 4.06e-01 |
| | $\alpha$ | 1.07e+01 | 1.92e-04 | 2.81e-02 | 1.22e-04 | 1.43e-01 |
| | $\beta$ | 6.90e+00 | 2.73e-03 | 1.68e-01 | 1.81e-03 | 1.62e-01 |
| Cls 3 | $\theta$ | 2.05e+00 | 1.44e-01 | — | — | — |
| | $\alpha$ | 2.29e+00 | 1.16e-01 | — | — | — |
| | $\beta$ | 1.78e+00 | 1.82e-01 | — | — | — |
| Cls 4 | $\theta$ | 3.82e+00 | 2.93e-02 | 3.73e-01 | 2.20e-02 | 3.45e-01 |
| | $\alpha$ | 9.61e+00 | 3.34e-04 | 4.70e-02 | 2.08e-04 | 1.44e-01 |
| | $\beta$ | 8.75e+00 | 6.15e-04 | 2.59e-01 | 4.26e-04 | 3.68e-02 |
| Cls 7 | $\theta$ | 4.20e+00 | 2.13e-02 | 4.03e-02 | 4.18e-02 | 1.00e+00 |
| | $\alpha$ | 1.15e+01 | 9.21e-05 | 8.56e-04 | 2.21e-04 | 9.02e-01 |
| | $\beta$ | 2.48e+01 | 5.47e-08 | 1.39e-06 | 3.37e-07 | 9.10e-01 |
| Cls 9 | $\theta$ | 1.69e+00 | 2.04e-01 | — | — | — |
| | $\alpha$ | 4.41e+00 | 2.20e-02 | 4.53e-01 | 1.71e-02 | 2.11e-01 |
| | $\beta$ | 3.09e+00 | 6.20e-02 | 7.41e-01 | 5.61e-02 | 2.30e-01 |

**Figure 10.**
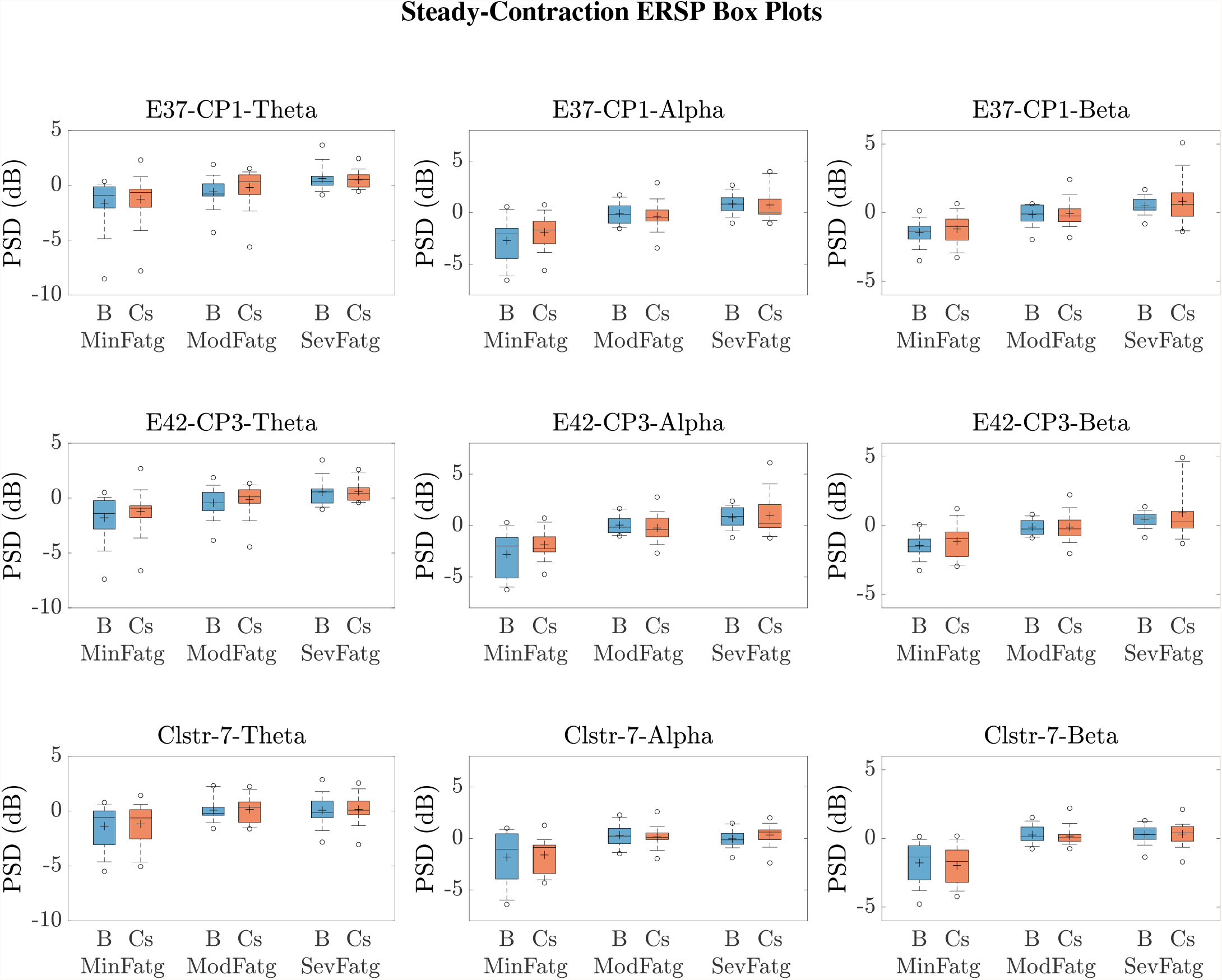
To study the modulation of band-specific ERSP values across the fatigue conditions, the time-averaged PSD values for the steady contraction and baseline were averaged over the EEG frequency bands of interest. The box plots of 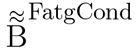 and 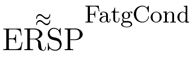 in the theta, alpha, and beta frequency band for CP1 (row 1) and CP3 (row 2) affirm that the theta-, alpha-, and beta-band PSD scale with the fatigue level during the baseline (B) and steady-contraction (Cs) interval. Likewise, the theta-, alpha-, and beta-band PSD computed for IC Cluster 7 proportionally vary with fatigue during B and Cs period (row 3).

**Figure 11.**
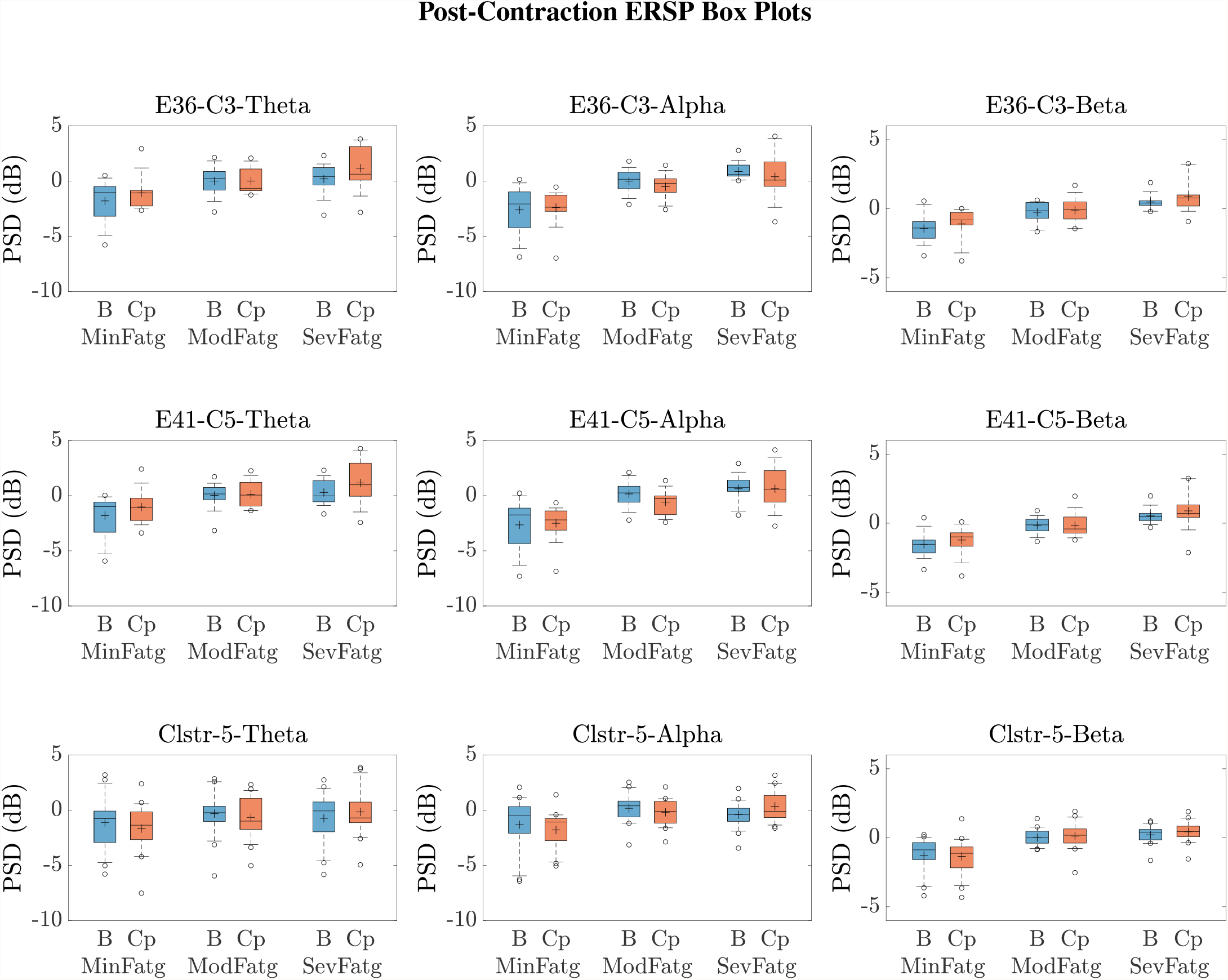
Given a frequency band of interest, the PSD values were averaged over the post-contraction and the respective baseline interval. From the box plots of 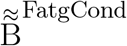 and 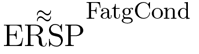 for the theta, alpha, and beta frequency band, one can infer that the band-specific time-averaged PSD scales with fatigue for C3 (row 1), C5 (row 2), and IC Cluster 5 (row 3) during the baseline (B) and post-contraction (Cp) period.

The subject-wise baseline PSD averaged over the trials in a given fatigue condition 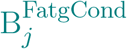 was log-transformed as 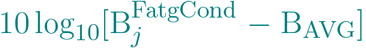. The quantity thus expressed in dB was averaged over the task-related (steady or post-contraction) baseline time interval and a specific frequency band, denoted as 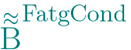 collectively for *N* subjects (components). In Fig. 10 and 11, the box plot of band-specific 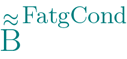 B for a fatigue condition is displayed beside the respective box plot of 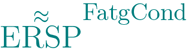 for the steady contraction and post-contraction, respectively. In IMT-related EEG channels and IC clusters, 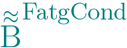 increases as the fatigue grows in concordance with the changes observed in 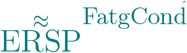 across the fatigue conditions. A reasonable explanation could be that the fatigue task affects the brain activity at rest in the same manner as during the steady or post-contraction, suggesting either a compensatory mechanism to reduce inhibition for facilitating motor planning and execution or a direct manifestation of neural fatigue. Recall from Section 4 that a common baseline B_AVG_ is derived using (4) in our analysis in order to counteract the changes induced by fatigue in the baseline. To summarize, 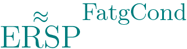 as well as 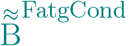 remain proportional to the fatigue level as portrayed in Fig. 10 and 11.

A critical aspect of this analysis stems from the fact that human EEG signals are inherently non-stationary due to noise and voluntary neural dynamics (Yang et al., 2021; Raza et al., 2019). In the current study design, since the ERSP representing MinFatg, ModFatg, and SevFatg state is estimated from EEG data acquired chronologically in time, it is highly probable that the fatigue-induced changes in ERSP overlap the signal non-stationarities. Furthermore, as shown in Fig. 10 and 11,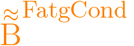 exhibits a trend similar to 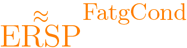 as fatigue progresses. In order to empirically tease out the confounding factor owing to the non-stationarity of EEG signals, we repeated the same analysis with motor/fatigue-unrelated channels and IC sources to inspect if the ERSP-fatigue proportional relationship still persists. In the strictest sense, 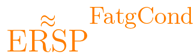 computed for the brain sources dissociated from motor areas is not expected to scale with fatigue, given that the results reported are noncontingent on EEG signal non-stationarities. Interestingly enough, as exemplified in Fig. 12, the brain sources not confined to motor-related cortical areas did fail to maintain the monotonically increasing trend with growing fatigue, thus mitigating this dilemma. Nevertheless, the discussion on why the ERSP computed for unrelated channels and neural sources in practice may still carry the traces of fatigue-induced changes is deferred to Section 7.

**Figure 12.**
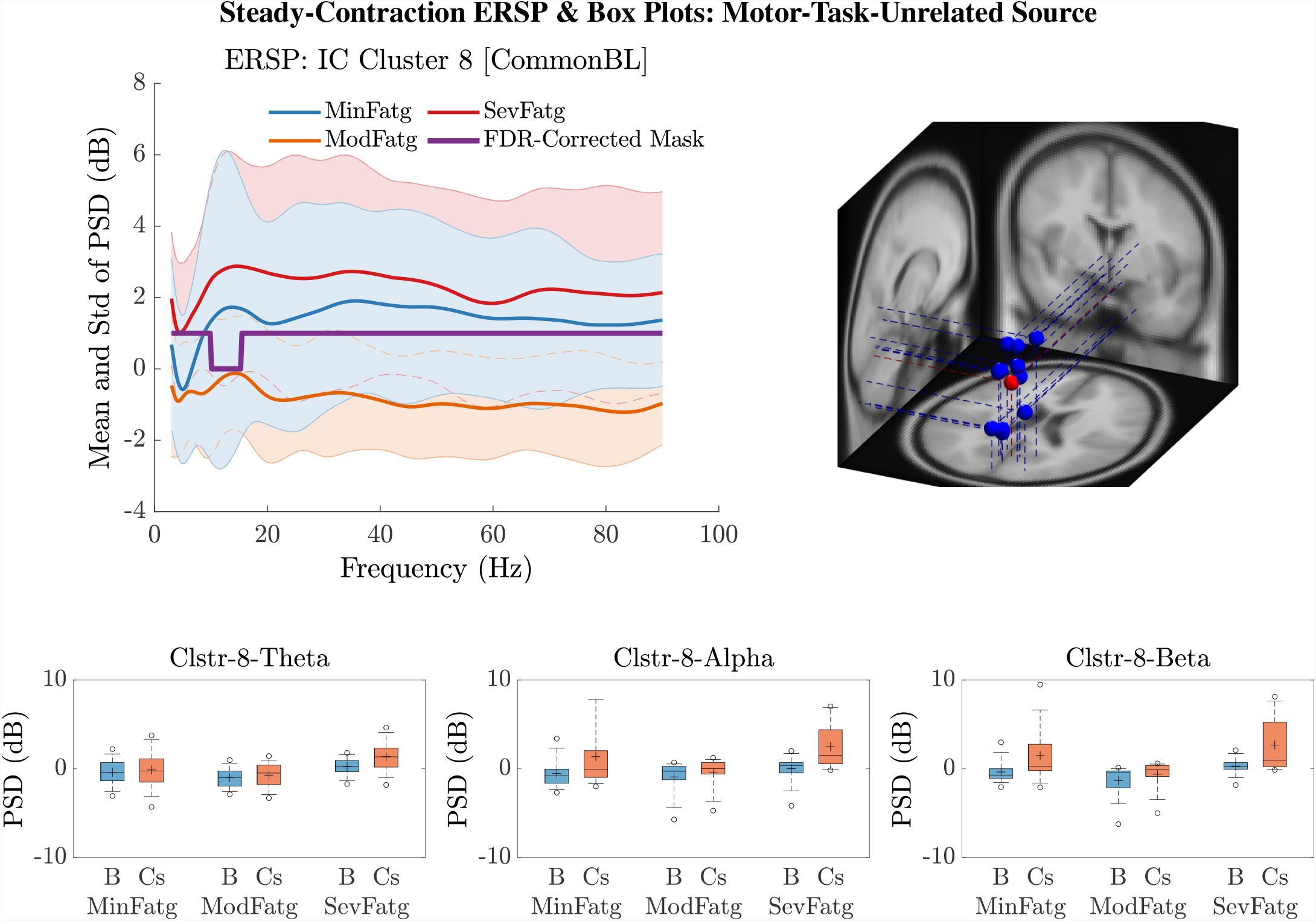
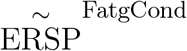 for IC Cluster 8 computed with the steady-contraction EEG data bears evidence to the fact that the time-averaged PSD fails to scale with the fatigue level throughout the frequency range of interest (row 1: left panel) since the brain source is not located inside a motor-related cortical area (row 1: right panel). The time-averaged band-specific PSD remains disproportional to the fatigue level as can be noted from the box plots of 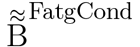 and 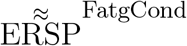 ERSP for the theta, alpha, and beta frequency band (row 2).

The *F* - and *p*-values quantifying the statistical significance of pairwise ERSP differences between the fatigue groups—Min-Mod Fatg, Min-Sev Fatg, and Mod-Sev Fatg—for theta, alpha, and beta band are reported in Table 1 (steady contraction) and Table 2 (post-contraction). We remark that a majority of pairwise differences between MinFatg and SevFatg remain statistically significant (*p* < 0.05 and *p* < 0.001). Even though the rest of the ERSP differences—MinFatg vs. ModFatg and ModFatg vs. SevFatg—are only seldom marginally significant (*p* < 0.1) supposedly due to a large inter-subject variability, the ERSP-fatigue proportional relationship is well preserved. Based on this premise, we conclude that during the steady as well as post-contraction, as fatigue progresses due to a prolonged IMT involving upper extremities, the PSD estimates from EEG channels and IC clusters associated with sensory and motor cortex have the tendency to increase in proportion to the fatigue level.

**Table 2.** The one-way ANOVA test and the post-hoc analysis (if *p* < 0.05 from ANOVA) resulted in the following *F* - and *p*-values for the time-averaged theta-, alpha-, and beta-band 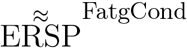 computed from the EEG signals and IC sources during post-contraction. The light, medium, and dark shades of blue mark the increase in PSD as fatigue escalates with a statistical significance of *p* < 0.1, *p* < 0.05, and *p* < 0.001, respectively.

| Channel/<br>IC Cluster | Frequency<br>Band | ANOVA<br>$F$-Value | ANOVA<br>$p$-Value | Min-Mod Fatg<br>$p$-Value | Min-Sev Fatg<br>$p$-Value | Mod-Sev Fatg<br>$p$-Value |
| --- | --- | --- | --- | --- | --- | --- |
| FC3 | $\theta$ | 5.91e+00 | 5.74e-03 | 1.84e-01 | 3.96e-03 | 2.41e-01 |
| | $\alpha$ | 8.43e+00 | 9.06e-04 | 1.88e-02 | 8.18e-04 | 4.97e-01 |
| | $\beta$ | 7.94e+00 | 1.28e-03 | 1.39e-01 | 8.18e-04 | 1.17e-01 |
| C1 | $\theta$ | 7.44e+00 | 1.83e-03 | 1.48e-01 | 1.19e-03 | 1.40e-01 |
| | $\alpha$ | 1.17e+01 | 1.08e-04 | 6.30e-03 | 9.00e-05 | 3.28e-01 |
| | $\beta$ | 8.71e+00 | 7.48e-04 | 4.65e-02 | 5.05e-04 | 2.26e-01 |
| C3 | $\theta$ | 7.19e+00 | 2.20e-03 | 1.74e-01 | 1.45e-03 | 1.35e-01 |
| | $\alpha$ | 1.05e+01 | 2.23e-04 | 1.16e-02 | 1.76e-04 | 3.21e-01 |
| | $\beta$ | 1.20e+01 | 8.93e-05 | 4.48e-02 | 5.19e-05 | 5.37e-02 |
| C5 | $\theta$ | 6.43e+00 | 3.87e-03 | 1.51e-01 | 2.62e-03 | 2.26e-01 |
| | $\alpha$ | 1.42e+01 | 2.31e-05 | 6.71e-03 | 1.49e-05 | 1.15e-01 |
| | $\beta$ | 1.29e+01 | 4.96e-05 | 4.29e-02 | 2.83e-05 | 3.60e-02 |
| CP1 | $\theta$ | 1.21e+01 | 8.23e-05 | 6.22e-02 | 4.79e-05 | 3.62e-02 |
| | $\alpha$ | 1.14e+01 | 1.25e-04 | 9.83e-03 | 9.38e-05 | 2.58e-01 |
| | $\beta$ | 9.00e+00 | 6.09e-04 | 5.71e-02 | 3.91e-04 | 1.65e-01 |
| CP3 | $\theta$ | 8.94e+00 | 6.36e-04 | 4.56e-02 | 4.23e-04 | 2.07e-01 |
| | $\alpha$ | 1.19e+01 | 9.08e-05 | 7.21e-03 | 7.05e-05 | 2.68e-01 |
| | $\beta$ | 1.28e+01 | 5.35e-05 | 2.95e-02 | 3.08e-05 | 5.52e-02 |
| Cls 4 | $\theta$ | 2.27e+00 | 1.12e-01 | — | — | — |
| | $\alpha$ | 9.40e+00 | 2.80e-04 | 1.42e-02 | 2.27e-04 | 3.77e-01 |
| | $\beta$ | 9.13e+00 | 3.47e-04 | 5.76e-02 | 2.10e-04 | 1.41e-01 |
| Cls 5 | $\theta$ | 2.61e+00 | 8.25e-02 | — | — | — |
| | $\alpha$ | 1.17e+01 | 6.22e-05 | 3.00e-03 | 6.53e-05 | 4.68e-01 |
| | $\beta$ | 1.60e+01 | 3.59e-06 | 1.67e-04 | 6.84e-06 | 6.31e-01 |
| Cls 6 | $\theta$ | 2.36e+00 | 1.10e-01 | — | — | — |
| | $\alpha$ | 7.96e+00 | 1.51e-03 | 6.47e-02 | 1.04e-03 | 2.46e-01 |
| | $\beta$ | 7.99e+00 | 1.48e-03 | 1.55e-01 | 9.69e-04 | 1.06e-01 |
| Cls 7 | $\theta$ | 8.78e+00 | 7.84e-04 | 1.28e-03 | 5.42e-03 | 8.62e-01 |
| | $\alpha$ | 9.52e+00 | 4.80e-04 | 5.57e-03 | 6.33e-04 | 7.20e-01 |
| | $\beta$ | 6.27e+00 | 4.61e-03 | 2.67e-02 | 5.64e-03 | 8.15e-01 |

### 6.5 Can ERSP Help Classify Fatigue Levels

As reported earlier, the ERSP pertaining to a fatigue condition exhibits a large inter-subject variability, which would pose challenges if ERSP is regarded as a standalone biomarker or feature to detect the level of fatigue within a multi-class classification framework. To further explore, we attempted to classify 12-D ERSP feature vectors deduced from the steady- and post-contraction EEG data into MinFatg, ModFatg, and SevFatg with a linear support vector machine (SVM). Each feature vector comprises the ERSP from individual trials computed for six EEG channels (FC3, C1, C3, C5, CP1, and CP3) and two frequency bands (alpha and beta) of interest. The accuracy (Ac) of binary classification (MinFatg and SevFatg) from a 10-fold cross validation is 78.22% for the steady-contraction and 71.72% for the post-contraction EEG data. When all the three classes were taken into account, as anticipated, Ac deteriorated to 55.78% and 55.23% for the data acquired during the steady- and post-contraction interval, respectively. The confusion matrices for the binary and three-class classification are presented in Fig. S3 (Supplementary Material). This means that the classifier could perform only marginally better than a chance-level assignment (33%) if three fatigue levels are considered. Interestingly, the classification results corroborate with the *p*-values from the post hoc analysis (recorded in Table 1 and 2), which reflect significant ERSP differences between MinFatg and SevFatg and not so between the remaining pairs—MinFatg vs. ModFatg and ModFatg vs. SevFatg. We remark that neither were the SVM hyperparameters tuned nor were an appropriate kernel function and the optimal feature set sought to enhance the classifier performance. In conclusion, even though our findings underscore that the ERSP during the steady- and post-contraction period in the alpha and beta EEG frequency band scales with fatigue levels, thus serving as a reliable proxy for central fatigue, implementing a machine learning approach to accurately predict the fatigue level from ERSP values is not a trivial extension and hence beyond the scope of our study.

## 7 DISCUSSION

### Cortical Regions/Channels of Interest

For the ERSP analysis, we have selected the EEG signals recorded from the electrodes, namely FC3, C1, C2, C3, CP1, and CP3, that overlay the following brain regions: (i) motor cortex comprising M1 (Brodmann area 4), premotor cortex, and SMA (Brodmann area 6); (ii) primary somatosensory cortex (Brodmann area 1, 2, 3); and (iii) superior parietal lobule (Brodmann area 5) (Donoghue and Sanes, 1994; Rosenbaum, 2009). The motor cortex is responsible for executing motor activities, especially contralateral upper-extremity muscle contractions as well as learned motor sequences. The premotor and SMA offer sensory guidance of movement, control proximal and trunk muscles, and plan complex and coordinated motor actions. The primary somatosensory cortex is associated with sensory perception involving arms and hands and helps with motor learning. The superior parietal lobule plays a crucial role in planned movements, receives sensory inputs from peripheral sources, and coordinates fine motor skills (Donoghue and Sanes, 1994; Rosenbaum, 2009). Since all the 14 individuals are right-handed, the EEG electrodes on the motor areas of the left (contralateral) hemisphere were selected.

### Relevance of IC Cluster Locations to Motor Task/Fatigue

The task-relevant IC clusters obtained from the steady and post-contraction EEG data point to neural sources residing in the following four regions: left cingulate gyrus, left superior parietal gyrus/left precuneus, right paracentral lobule (SMA), and left transverse temporal gyrus. These brain regions were found to have an association with the upper-extremity motor task performance and/or fatigue by the previous studies. The dorsal anterior cingulate cortex is responsible for the mediation of task-related motor control and has connections to premotor and sensorimotor areas (Asemi et al., 2015). In an fMRI study on the submaximal fatigue task, a steady increase of the BOLD signal in the cingulate gyrus has been observed during both the sustained and intermittent muscle contractions (Liu et al., 2003). The superior parietal cortex has been demonstrated to play a crucial role in the reach-to-grasp action that involves the elbow flexion movement (Cavina-Pratesi et al., 2010). An MEG study with the bimanual load-lifting task showed the involvement of precuneus, which is contralateral to the load-lifting forearm, in the elbow-joint flexion movement (Di Rienzo et al., 2019). The SMA is an important source of upstream drive to M1 and sends projections to M1 (Muakkassa and Strick, 1979). It has an extensive transcallosal connectivity and plays a critical role in uni/bimanual muscle activation (Grefkes et al., 2008). Reportedly, the fatiguing tasks reduce the SMA activity, which in turn decrease the excitatory drive to the corticospinal neuron pool (Sharples et al., 2016).

### Frequency Bands of Interest

We investigated the ERSP measure computed in the theta, alpha, and beta band across three fatigue conditions for the following rationale. **Theta Frequency Band**. An increase in the region-specific theta-band activity has been observed with sensorimotor processing (Cruikshank et al., 2012). The theta activity on the contralateral motor cortex is modulated by acceleration and associated with adaptive adjustments during upper limb movements (Ofori et al., 2015). The theta-band power in the contralateral motor cortex serves as a marker for enhanced control engagement across cortical regions. Moreover, theta synchrony observed at the start of movement in the premotor cortex and anterior cingulate cortex is critical for motor execution (Ofori et al., 2015; Paus, 2001). Therefore, the theta-band power is thought to be linked to the upper limb movement and the acceleration of the movement (Ofori et al., 2015). **Alpha Frequency Band**. A decrease in the alpha-band activity is commonly encountered during movement [referred to as event-related desynchronization (ERD)], which reflects the excitability of neurons due to activation in a specific cortical region (Toro et al., 1994; Babiloni et al., 1999; Neuper and Pfurtscheller, 2001). A suppression in the alpha-band activity is interpreted as a release from the inhibition caused by alpha ERD (Klimesch, 2012). This phenomenon has been observed in the ipsilateral motor and somatosensory areas near the movement cessation in proportion to the task demands (Ofori et al., 2015). **Beta Frequency Band**. The beta-band spectral power tends to reduce over bilateral motor cortex at the beginning of movement (Cruikshank et al., 2012; Gwin and Ferris, 2012; Pastötter et al., 2012; Kilavik et al., 2013). However, the beta-band activity in the contralateral motor area is not altered at the acceleration phase of movement but it increased during the deceleration phase (Ofori et al., 2015).

### fMRI Studies with Corroborating Findings

The study findings corroborate with the inferences drawn from many fatigue studies. Imaging studies conducted during fatigue tasks involving single-limb contractions induced changes in the activity of motor cortex and other brain regions. Engaging in prolonged submaximal contractions with dominant hand muscles progressively increased the fMRI BOLD signal from the contralateral and ipsilateral sensorimotor cortex, SMA, and cingulate motor cortex (Liu et al., 2003; Van Duinen et al., 2007; Benwell et al., 2007). The activation in the motor cortex (Post et al., 2009) and sensorimotor cortex (Liu et al., 2002; Steens et al., 2012) increased during sustained maximal efforts as inferred from the BOLD signal. The increase in the BOLD signal due to submaximal and maximal tasks is attributed to simultaneous excitatory and inhibitory activity, recruitment of neurons to engage muscles noncritical to the task, and effects induced by the modified sensory feedback (Post et al., 2009). However, in the case of submaximal efforts, an augmented BOLD signal implies an increase in motor cortex output due to excitatory input (Taylor et al., 2016). In a fatigue task study involving repetitive MVCs of finger flexor muscles (Liu et al., 2005b), the fMRI signal level in the primary, secondary, and association motor-function cortices remained unaltered throughout the task as opposed to the findings from a continuous muscle contraction (Yang et al., 2010). The fMRI studies suggest that disparate cell populations might be responsible for controlling sustained and intermittent voluntary motor activities. Thus, our study offered additional evidence to support that the motor cortical centers control the repetitive and continuous fatigue task differently.

### MEG Studies

An MEG-based study concerning the influence of physical fatigue due to submaximal contractions on the PMBR and movement-related beta decrease (MRBD) within the contralateral sensorimotor cortex concluded the following: the physical fatigue gives rise to (i) an enhanced PMBR and (ii) cancellation of attenuation in MRBD related to task habituation (Fry et al., 2017). The MEG beta- and gamma-band power reduction with respect to relaxation period was less pronounced during the execution of a non-fatiguing motor task performed after producing muscle fatigue (Tecchio et al., 2006). In contrast to our findings, a fatigue-inducing task session with repetitive handgrips at MVC led to a decrease in the MEG alpha-band power in the ipsilateral sensorimotor and prefrontal areas (Tanaka et al., 2015).

### EEG Studies with Corroborating Findings

Submaximal handgrip isometric contractions inducing progressive fatigue resulted in a significant increase in MRCP at the precentral (Cz and FCz) and central contralateral (C3) electrode sites (Johnston et al., 2001). In like manner, the central fatigue induced by submaximal (Guo et al., 2014) and lower limb (Berchicci et al., 2013) isometric contractions increased the amplitude of MRCP in M1, prefrontal cortex, SMA, and premotor cortex. The muscle fatigue was reported to increase the MRCP amplitude during movement execution (De Morree et al., 2012). A significant increase in EEG power within the alpha and beta band was noticed as fatigue developed from a high-intensity cycling exercise in the following cortical regions: parietal and limbic regions (Schneider et al., 2009); Brodmann area 11 (both alpha and beta bands) and parietal lobe (only beta band) (Hilty et al., 2011); SMA, frontal, and parietal lobes (Enders et al., 2016). During repetitive force grip submaximal contractions, the area under the curve of readiness potential^4^ doubled at electrode Cz and increased fourfold at electrodes C3’ and C4’ (1 cm anterior to C3 and C4, respectively) that cover the motor cortex (Schillings et al., 2006).

### EEG Studies with Contradictory Findings

Contrary to what we have observed in the submaximal contractions, the intermittent handgrip MVCs were reported to have significantly decreased the MRCP with fatigue during the sustained phase of muscle contractions. A significant fatigue-related EEG power decline was observed at alpha 2 (C3), beta 1 (C3 and Fz), and beta 2 (C3, Cz, and C4) bands (Liu et al., 2005a). Similar contradictory outcome was reported from a sustained (submaximal) voluntary handgrip contraction to fatigue (Nishihira et al., 1995); a decrease in alpha 1 and alpha 2 power was observed at Cz, C4, P3, and P4 during the second half of the contraction compared to the first half, whereas the power increased at the frontal position. Moreover, the mean EEG power measured during the rest, pre-fatigue, and post-fatigue state from a sustained Adductor pinch task until fatigue performed at two different target values did not produce a significant difference across the three states in alpha, beta, or gamma band (Sh et al., 2020).

### Influence of Age and Gender on the PSD-Fatigue Relationship

Even though the participants’ age falls within the range 26−72 years, only two of them exceeded 65 years. This means that the rest of the subjects were middle-aged (26−59 years). Therefore, we refrain from comparing the fatigue-related PSD modulation across different groups based on the age criterion, e.g., two groups consisting of young (< 65 years) and old (*≥* 65 years) participants. Nevertheless, we repeated the analyses by excluding the EEG signals from older (*≥* 65 years) subjects to verify whether the results could have been impacted by the data from the two outliers—subjects of age 70 and 72 years—in the former analyses. Interestingly, the outcome of both the channel- and source-level-analysis with the data from the remaining 12 subjects is in total agreement with what we have already inferred from the original dataset. Recall that the dataset consists of EEG from six male and eight female volunteers. To make sure that the conclusions arrived at are gender-independent, we carried out the channel-level analysis with the male and female participants separately and examined the outcome. Notwithstanding the gender difference, the spectral power varies relative to the fatigue level for the data from the channels of interest, provided theta-, alpha-, and beta-band frequencies are considered for statistical inferences (refer to Fig. S4, Supplementary Material).

### Future Perspectives

Past studies have reported the association of fatigue with the following frequency bands in pathological conditions: theta, alpha, or beta band in chronic fatigue syndrome (Siemionow et al., 2004; Flor-Henry et al., 2010) and multiple sclerosis (Leocani et al., 2001; Vecchio et al., 2017); alpha and/or beta band in burnout syndrome (Luijtelaar et al., 2010); theta, alpha, beta, and gamma band in cancer (Allexandre et al., 2020). In this respect, our study outcome will provide a reference and help investigate how fatigue would alter the spectral power in pathologies in comparison to healthy individuals. In particular, our future research direction would be to investigate how the EEG-based spectral power changes due to fatigue compare between cancer or multiple sclerosis patients and healthy volunteers, thus enhancing our understanding of the effect of a brain pathology on the central fatigue mechanism.

Furthermore, in lieu of the common baseline considered in this work that takes into account the rest period corresponding to all the trials, the common baseline derived from only the MinFatg trials may be used. We tested both the baseline options and the findings were consistent.

### Study Limitations

Although the EEG data represent brain activities related to fatigue conditions derived from the scalp electrodes and underlying source regions, the data are influenced by connectivity with other cortical and subcortical areas (Jiang et al., 2012, 2016), which could not be determined by this study. Aside from a small sample size, a major limitation of this study is the inter-subject variability in terms of the ERSP measure. For instance, 1-D ERSP plots from two subjects are portrayed in Fig. S5 (Supplementary Material), where the PSD changes with respect to fatigue levels within a frequency band of interest are quite dissimilar. Admittedly, in a few subjects as exemplified in these plots, we failed to notice a monotonous increase of PSD in a specific motor-related cortical area (or EEG channel) within the theta, alpha, or beta frequency band as the perceived fatigue increases. One plausible explanation could be that the availability of multiple cortical centers that control one motor task or muscle group makes it possible to “ shift the activation center” to compensate for fatigue as demonstrated in (Liu et al., 2007). However, this phenomenon was not widely observed and the avoidance of central fatigue by shifting neuron populations linked to a fatiguing motor task requires further investigation.

In the channel-level ERSP analysis, the EEG recordings do not entirely represent the electrical potential generated by the cortical tissues right beneath the respective scalp electrodes due to the volume-conduction effect (Nunez et al., 1997). The scalp EEG signals will further be linearly combined when they are reconstructed with the neural ICs extracted via IC pruning. On the other hand, the source-level analysis is carried out with IC clusters produced by *k*-means algorithm, which does not guarantee that the ICs in each cluster are in close proximity to each other and located in one functional area of cerebral cortex. Furthermore, the source localization is performed with the DIPFIT toolbox in EEGLAB, which introduces inaccuracies—deviation between the location of estimated dipoles and true sources—in the order of around 10 mm (Knyazev, 2013). Also, the lack of subject-specific MRIs and 3-D-digitized EEG electrode locations in the present study would interfere with the precise estimation of dipole positions. The aforementioned factors would have led to the overlapping of information borne by different scalp electrodes or IC clusters in the channel- or source-level analysis, respectively. Consequently, the traces of fatigue-induced alterations in ERSP are likely to be present even in channels and sources that are not directly linked to motor tasks.

### Pointers for Improvement

The inherent shortcomings of EEG can be overcome by performing the analysis with other neuroimaging techniques having a better spatial resolution, e.g., electrocorticography and fMRI. To enhance the source localization accuracy, the future studies may investigate distributed source modeling methods recommended in Handiru et al. (2018) as well as incorporate individual MRIs and 3-D-digitized EEG positions (Shirazi and Huang, 2019).

We remark that the ERD—decrease of cortical spectral power in the alpha and beta frequency bands during contractions in relation to the baseline period—was not conspicuous, which is contrary to the observation reported in (Cremoux et al., 2013) with the EEG data from an isometric elbow flexion task at different force levels. We speculate that it could be because of our experimental design dedicated to study the effect of fatigue on the cortical spectral power during the steady^5^- and post-contraction period. Finally, we opine that the source-level analysis can be further improved by employing more sophisticated source-separation approaches instead of traditional ones that assume that the underlying sources have a large kurtosis, whereas the distributions of EEG/MEG sources tend to be multimodal (Knuth, 1998).

## 8 CONCLUSIONS

In the present study, by dividing the EEG data from intermittent submaximal elbow flexions until subjective exhaustion into three equal segments, each one representing progressive levels of fatigue—minimum, moderate, and severe—the changes in various EEG band power with regard to fatigue levels were studied. **Key Findings in a Nutshell**. Our channel- and source-level analyses offer insights into the effect of progressive fatigue during a prolonged intermittent contraction task on the EEG cortical rhythms: (i) The theta, alpha, and beta PSDs scale with fatigue in cortical motor areas. These signal changes most likely are associated with progressively increasing the effort to continue the motor task to compensate for progressive fatigue. The fMRI (Dai et al., 2001) and EEG-measured MRCP (Siemionow et al., 2000) studies have reported a linear relationship between fMRI/MRCP signals and level of effort (level of voluntary force). (ii) The pairwise PSD differences between the fatigue conditions maintain a statistical significance—especially in minimum vs. severe fatigue. (iii) The relationship described in (i) and (ii) between the band-specific spectral power and the fatigue level is exhibited by the steady- and post-contraction EEG data. (iv) The source-level analysis reveals that the fatigue effect on PSD is prevalent in SMA, superior parietal cortex, and cingulate gyrus during the steady and post-contraction period. (v) The age-related and gender-based differences do not seem to have an influence on the fatigue-PSD behavior.

To the best of our knowledge, the spectral power changes in relation to graded fatigue levels were not investigated by past studies. Furthermore, as mentioned earlier, the inferences drawn would help clarify the inconsistencies and contradictions among the findings from fMRI, MEG, and EEG fatigue studies involving intermittent/sustained maximal/submaximal contractions. Importantly, since the results point to fatigue-induced changes in the cortical activation in healthy individuals, they would serve as a reference to evaluate the alterations in spectral power caused by fatigue in pathological conditions such as cancer and multiple sclerosis. We envisage that the neural underpinnings of graded fatigue would provide pointers to design effective rehabilitation therapies for patients suffering from movement disorders and related fatigue symptoms.

## AUTHOR CONTRIBUTIONS

SES led the research study in terms of conceptualization, scripting, data analysis, interpretation of results, and manuscript writing. VSH was involved in conceptualization, scripting, data analysis, interpretation of results, and writing. DA assisted in data analysis, interpreting the results, and writing. AH preprocessed the data. SS contributed to the research idea through discussion. GY oversaw the data collection and helped with conceptualization, interpretation of results, and writing. All authors contributed to the article and approved the submitted version.

## ACKNOWLEDGMENTS

The authors would like to thank Shyamala Magdalene for her help with the illustrations and proofreading. They also thank the anonymous reviewers for their constructive comments that helped significantly improve the earlier version of the article. The first author gratefully acknowledges Prof. Umberto Amato, CNR, Italy, for his insightful suggestions on the statistical framework followed in this work.

## SUPPLEMENTAL DATA

The supplementary material consists of additional study results presented in the discussion section.

## DATA AVAILABILITY STATEMENT

The raw EEG and force data are not publicly available due to IRB restrictions. However, the MATLAB scripts and the de-identified preprocessed EEG data associated with this study can be shared, if requests are made to the corresponding author.

## Footnotes

1 Readers are cautioned not to confuse this abbreviation with the conventional one for event-related synchronization.

2 To obtain physiologically meaningful interpretation of results, both the channel-level and source-level ERSP were computed for individual EEG frequency bands as each one has different functional characteristics. The typical frequency bands and their approximate spectral boundaries in adults (Tatum IV, 2021) are as follows: delta - 1 Hz to 4 Hz; theta - 4 Hz to 8 Hz; alpha - 8 Hz to 13 Hz; beta - 13 Hz to 30 Hz; low gamma - 30 Hz to 50 Hz; and high gamma - 50 Hz to 80 Hz.

3 An IC cluster index is meant only for referring to a particular IC cluster without any other significance.

4 A negative movement-related cortical EEG potential appearing over the scalp about 1 s prior to a self-paced motor act.

5 This duration is 1 s to 3 s after the task onset, whereas the ERD should attain the peak value right after the onset.

