## Supplementary Material for "EEG-Based Spectral Analysis Showing Brainwave Changes Related to Modulating Progressive Fatigue During a Prolonged Intermittent Motor Task"

### Review Response 1: Aricle Resubmitted to Front. Hum. Neurosci. - Brain Imaging and Stimulation

January 19, 2022

Please find below the response to the review comments received concerning our manuscript entitled, "EEG-Based Spectral Analysis Showing Brainwave Changes Related to Modulating Progressive Fatigue During a Prolonged Intermittent Motor Task," submitted to Front. Hum. Neurosci. Journal under the research topic "The Brain Under Fatigue" for possible consideration of the same as an original research article.

#### Response to the Editorial Team

At the outset, the authors gratefully acknowledge the review efforts by the editor and reviewers towards handling the manuscript. Primarily, they would like to thank the reviewers for their constructive and insightful comments, which enabled them to provide a few more essential details, include additional results, and rewrite some portions to enhance clarity and understanding.

Please note that while revising the article, we realized that there is an elegant way of performing the statistical analysis. Therefore, we adopted the changes owing to the improved statistical design. Consequently, the results reported in the revised version are physiologically interpretable and more meaningful. This additional work induced further delay in the revision process. We, per the suggestion of Reviewer 1, consulted with a senior author to ensure the clarity of writing and a statistical expert (Prof. Umberto Amato, CNR, Italy, whom we acknowledged in the manuscript) to verify the correctness of the statistical design.

Following each comment of a reviewer in red, the response is stated in black, and where it is addressed in the manuscript is mentioned in blue. In addition, the modifications in the revised manuscript to account for the comments of Reviewer 1 and Reviewer 2 are in violet and teal, respectively; following the reviewers' approval, they will be turned into black.

#### Response to Reviewer 1

We are indebted to express our sincere gratitude to Reviewer 1 for his meticulous list of valuable suggestions for further improvement. We have carefully considered all the comments while revising the paper. Mentioned below are the changes made in the revised version.

##### Specific Comments

- At the end of Introduction section, the author mentioned "The secondary goals are to ... by a pathology and to design effective rehabilitation techniques targeting patients ... ". Is this the goal for the current study or a future study? Please clarify.

We admit that it is a future goal, and hence corrected the sentences in the revised version.

The corrected sentences are on p. 4, §1, para. 2, ll. 9-13.

- I suggest adding a table describing the basic information of subjects such as age, sex, P values, ...

  We agree with this suggestion and included this information. Since we do not have more than one group, there is no $p$-value to report. Instead, we listed relevant demographic details—age, gender, BMI (body mass index), endurance, and BFI (brief fatigue inventory).

  Kindly refer to p. 4, §2.1, last sentence. For want of space, the table is provided on p. 1, Supplementary Material. If the reviewer feels that it is better to place Table S1 in the main manuscript, we will do so.

- What is the reason to employ 40% MVC?

  The rationale for selecting the effort level as 40% MVC is explicitly stated in the revised version.

  Please refer to p. 5, §2.2.2, ll. 2-5.

- What is ERSP? Please define ERSP.

  We have included both an informal and formal definition of ERSP.

  We have introduced the term ERSP informally on p. 3, §1, para. 1, ll. 4-7. A more rigorous definition is provided on pp. 8 & 9, §4, eqn. (2) & (4), and in the associated text.

- What is the reason to employ the default parameters for $k$-means? How do you know the default parameters fit your dataset and the current study's hypothesis?

  The reason for the default choice of $k$-means parameters (especially the number of clusters) and its suitability to the present dataset are described.

  The parameter choice in $k$-means is explained on p. 6, §4, para. 1, ll. 8-11.

- There are over-detailed descriptions in this manuscript. For example, you do not need to write several lines to introduce what ANOVA or FDR are. I am also not sure why the author placed figure 2 and 4 in this manuscript. Also, Figure 5, I can hardly understand what this figure represents or what it wants to show. There are so many other contents that are not necessary. it will be very beneficial to have a senior author carefully revise the entire manuscript to make it more readable.

  We addressed this major concern in the revised manuscript as follows.

  The over-detailed descriptions related to §5 (ANOVA and FDR) are carefully edited or removed. Consequently, §5 (pp. 9 & 10) is considerably shortened. We moved Fig. 2 & 4 (initial version) to Supplementary Material, thereby the readers will not get distracted by too many illustrations. To better understand §3, we have rewritten some portions and organized the preprocessing steps under suitable subheadings. We have extensively altered the presentation of **Statistical Framework** in §4 (pp. 8 & 9) to enhance the readability and corrected some notations. A subfigure in Fig. 5 (earlier version)—EEG Electrode Locations—is removed and the figure caption (Fig. 3 of resubmitted version) is revised to give a lucid explanation of the IC clusters that are displayed. As suggested, by consulting a senior author of this manuscript and with the help of a statistical expert, the contents of §6 have been substantially modified. This rework resulted in physiologically more meaningful and interpretable results. In addition, some portions of §7 are rewritten, e.g., **Study Limitations** and **Pointers for Improvement**.
